# APOE Genotype Influences on The Brain Metabolome of Aging Mice – Role for Mitochondrial Energetics in Mechanisms of Resilience in APOE2 Genotype

**DOI:** 10.1101/2025.02.25.640178

**Authors:** Kamil Borkowski, Nuanyi Liang, Na Zhao, Matthias Arnold, Kevin Huynh, Naama Karu, Siamak Mahmoudiandehkordi, Alexandra Kueider-Paisley, Takahisa Kanekiyo, Guojun Bu, Rima Kaddurah-Daouk, the Alzheimer’s Disease Metabolomics Consortium

## Abstract

Alzheimer’s disease (AD) risk and progression are significantly influenced by APOE genotype with APOE4 increasing and APOE2 decreasing susceptibility compared to APOE3. While the effect of those genotypes was extensively studied on blood metabolome, less is known about their impact in the brain. Here we investigated the impacts of APOE genotypes and aging on brain metabolic profiles across the lifespan, using human APOE-targeted replacement mice. Biocrates P180 targeted metabolomics platform was used to measure a broad range of metabolites probing various metabolic processes. In all genotypes investigated we report changes in acylcarnitines, biogenic amines, amino acids, phospholipids and sphingomyelins during aging. The decreased ratio of medium to long-chain acylcarnitine suggests a reduced level of fatty acid β-oxidation and thus the possibility of mitochondrial dysfunction as these animals age. Additionally, aging APOE2/2 mice had altered branch-chain amino acids (BCAA) profile and increased their downstream metabolite C5 acylcarnitine, indicating increased branched-chain amino acid utilization in TCA cycle and better energetic profile endowed by this protective genotype. We compared these results with human dorsolateral prefrontal cortex metabolomic data from the Religious Orders Study/Memory and Aging Project, and we found that the carriers of APOE2/3 genotype had lower markers of impaired BCAA katabolism, including tiglyl carnitine, methylmalonate and 3-methylglutaconate. In summary, these results suggest a potential involvement of the APOE2 genotype in BCAA utilization in the TCA cycle and nominate these humanized APOE mouse models for further study of APOE in AD, brain aging, and brain BCAA utilization for energy. We have previously shown lower plasma BCAA to be associated with incident dementia, and their higher levels in brain with AD pathology and cognitive impairment. Those findings together with our current results could potentially explain the AD-protective effect of APOE2 genotype by enabling higher utilization of BCAA for energy during the decline of fatty acid β-oxidation.

## 1. Introduction

APOE genotype, together with age, are major factors contributing to Alzheimer’s disease (AD), with APOE4 increasing and APOE2 decreasing the likelihood of AD development compared to the most common allele, APOE3 (1–3). The APOE genotypes are associated with the trajectory of brain amyloid and tau pathologies development (4, 5), brain structural atrophy (6, 7), and the severity of cognitive decline (8, 9). However, a comprehensive understanding of the metabolic pathways linking APOE genotypes and AD pathogenesis is still largely absent.

Metabolomics provides powerful tools to map dysregulation in metabolic processes related to diseases, including AD (10, 11). Alzheimer’s Disease Metabolomics Consortium (ADMC) has pioneered metabolomic profiling of large, well-characterized AD cohorts, including AD Neuroimaging Initiative (ADNI) and Religious Orders Study/Memory and Aging Project (ROS-MAP) and several animal models for AD, providing a comprehensive picture of central and peripheral metabolic alteration across AD trajectories (12–15). We have demonstrated a strong connection between peripheral and central metabolism and defined peripheral metabolic pathways and health markers that influence central processes in AD. Those include pathways related to amino acids (16) branch-chain amino acids (BCAA) (17), oxylipins and endocannabinoids (18, 19), ceramides and sphingolipids that pointed to an important function for S1P signaling (20), bile acids pointing to role for cholesterol clearance and gut microbiome function (21) among others (16, 17). In large studies including the Alzheimer’s Disease Neuroimaging Initiative (ADNI), the Australian Imaging, Biomarkers and Lifestyle (AIBL) and the Busselton Health Study (BHS) comprising of over six thousands individuals, we defined plasma lipid signatures for APOE genotypes, highlighting ether lipids as contributor to protective properties of APOE2 genotype (22). Furthermore, we identified impaired energy metabolism in APOE4 carriers (23).

Five decades of research link mitochondria to AD (24). Our own work led to development of a bioenergetic capacity index that enables patient stratification, highlighting individuals who have low capacity and could benefit from personalized approaches to improve mitochondrial function (25). Differences between the sexes and APOE genotypes in mitochondria energetics are also noted (10). The impaired β-oxidation can be assessed by the levels of fatty acids carnitines (AC), with the higher levels of long-chain AC being associated with AD, possibly involving compromised carnitine shuttle (26). Among other substrates for the TCA cycle, we have shown associations of lower levels of BCAA with incident dementia (17) and their higher levels in brain with AD pathology and cognitive impairment (27). Beyond association, a clinical trial including people with AD found that ketogenic diet improved some aspects of cognition, yet only in non-carriers of APOE4, pointing to the APOE-specific causative relationship between brain energetic metabolism and AD (28). Thus, understanding the involvement of APOE genotypes in the metabolism of the aging brain can help elucidate the disease mechanism and identify potential therapeutic targets for AD (29). Additionally, the connection between aging brain metabolism and AD is far from being understood.

Due to the low abundance of APOE2/2 and APOE4/4 heterozygotes in human studies and challenges in sampling the aging human brain, the utilization of animal models on human APOE background is an attractive alternative to human participants. Collective efforts are being made to construct useful animal models under MODEL-AD initiative (30), and the relevance of findings from these animal models to humans should be carefully evaluated. Our current work utilizes human APOE-targeted replacement (TR) mice (31) and the Biocrates P180 targeted metabolomics platform to investigate the influence of APOE2/2, APOE3/3 and APOE4/4 on metabolic trajectories of aging mice. The Biocrates P180 platform provides an insight into several areas of metabolism known to be affected in AD, including markers of fatty acid β-oxidation, amino acids, amines, phospholipids, and sphingomyelins (SMs) (32), with informative ratios and summations approximating enzymatic activities. Our Alzheimer Disease Metabolomics Consortium demonstrated utility of this platform for AD and aging – related biomarkers discovery (17). We have previously reported APOE genotype and age – related metabolic perturbations in serum, where APOE2/2 mice manifested greater levels of long-chain AC in older mice (33). In the current study, we describe the influence of APOE2/2, APOE3/3 and APOE4/4 on age-related metabolic trajectories of mice brain. Additionally, we used data from the Religious Orders Study/Memory and Aging Project (ROS-MAP), generated using the Metabolon^TM^ platform, to compare the relevance of the current mice model findings to humans.

## 2. Materials and Methods

### 2.1 Experimental mice

APOE-TR mice in which the murine Apoe gene is replaced with the human APOE2, APOE3, or APOE4 gene (31) were obtained from Taconic Biosciences. These animals were housed in environments with regulated temperature and lighting and were provided ad libitum access to food and water. The APOE-TR mice were harvested at 3, 12 and 24 months of age (n=8 mice/genotype/sex/age group) as described previously (33). Those time points correspond to adolescent; adult; old with the mice life expectancy of 26-30 months. The cortex tissues from the brain were collected and snap-frozen in liquid nitrogen, then stored at −80°C. All procedures involving animals received approval from the Mayo Clinic Institutional Animal Care and Use Committee and adhered strictly to the guidelines outlined in the National Institutes of Health Guide for the Care and Use of Laboratory Animals and Animal Research: Reporting of In Vitro Experiments.

### 2.2 Brain metabolites detection

The metabolites in these cortical samples were analyzed using the AbsoluteIDQ p180 kit (Biocrates Life Sciences AG, Innsbruck, Austria) according to the user manual. Briefly, weighed brain tissue samples were provided frozen in Precellys soft tissue homogenizing CK14 tubes (Bertin Technologies, Montigny-le-Bretonneux, France). Each sample was diluted with 100 μL 1:1 v/v methanol:water. Samples were then homogenized using three 10-second pulses in the Precellys Evolution between which samples were chilled using the Cryolys cooling system. After the first three cycles, 250 μL v/v 3:1 methanol:chloroform were added to each sample followed by three 10-second pulses between which samples were again chilled using the Cryolys cooling system. The homogenized samples were stored at −80C until the day of extraction with the p180 kits. On the day of p180 sample extraction the samples were thawed and vortexed. The samples were then centrifuged at 15000 rpm for 10 minutes in a refrigerated (4C) centrifuge then stored on ice until addition to the p180 kit plates. Samples were prepared using the AbsoluteIDQ® p180 kit (Biocrates Innsbruck, Austria) in accordance with their detailed protocol. After the addition of 10 μL of the supplied internal standard solution to each well of the 96-well extraction plate, 10 μL of each blank, calibration standard, and Biocrates QC samples were added to the appropriate wells. For the Brain SPQC and study samples 15 μL of each sample were added. The plate was then dried under a gentle stream of nitrogen for 10 minutes. An additional 15 μL of each SPQC and brain tissue homogenate sample were added to the appropriate wells followed by an additional 20 minutes of drying. The samples were derivatized with phenyl isothiocyanate then eluted with 5mM ammonium acetate in methanol. Samples were diluted with either 1:1 methanol:water for the UPLC analysis (4:1) or running solvent (a proprietary mixture provided by Biocrates) for flow injection analysis (20:1).

A pool of equal volumes of all 76 samples analyzed on the first plate was created (4920 Brain SPQC). The pooled sample was prepared and analyzed in the same way as the study samples on both plates. From each plate this sample was injected once before, once during, and once after the study samples to measure the performance of the assay across the sample cohort.

### 2.3 Data preprocessing

The raw metabolomics dataset had 182 metabolites for 144 samples. We first aligned measurement batches via sample-median quotient normalization. Next, we removed 21 metabolites that had significant amounts of missing data (>20% missing values). We then excluded one additional metabolite (citrulline) because it had a large coefficient of variation (0.359). Finally, metabolite concentrations were log2-transformed and remaining missing data was imputed using k-nearest neighbor imputation (using both sample and metabolite vectors with k=10). Multivariable analysis identified measurements for one 3-month-old APOE4 female mouse as probable sample outlier, which was subsequently removed from the analysis. The final metabolomics matrix used for analysis held data on 161 metabolites for 143 mouse samples.

### 2.4 Human data

Human dorsolateral prefrontal cortex (DLPFC) data of the ROS-MAP cohort (34), generated using the metabolomics platform from Metabolon Inc., were used to investigate the differences between the APOE2/3 (n=66), APOE3/3 (n=304) and APOE 3/4 (n=112) genotypes. The generation of metabolomic data for this analysis has been previously described (16). The demographics of the cohort are provided in Supplementary Table S1.

### 2.5 Statistical analysis

All statistical tests were performed using JMP Pro 16 (JMP, SAS Institute, Carry, NC). Prior to analysis, data were tested for outliers using the robust Huber M test, and missing data were imputed using multivariate normal imputation for variables which were at least 75% complete. Additionally, variables were normalized, centered, and scaled using Johnson’s transformation, with normality verification using the Shapiro–Wilk test. To facilitate interpretation, we reduced the dimensionality of the data using unsupervised variable clustering and generating a single cluster component value for variables within each cluster. Cluster components were generated using the JMP variable clustering algorithm, which uses the first principal component of the variables in that cluster. Clustering was performed separately for each metabolite chemical class. Factorial analysis was performed to evaluate the age, genotype, sex and the age x genotype interaction effects on the levels of brain metabolites. Additionally, analysis of covariance (ANCOVA) with contrast post-test was applied to determine the differences in means between each time point within each genotype and between the genotypes (sex was used as a covariate). The above analysis was performed on both cluster components and individual metabolites to ensure that the cluster components result represented the behavior of all metabolites within the cluster. For the phospholipids, the differences in the mean of each timepoint were analyzed using ANCOVA, with the genotype and sex as covariates and contrast post-test. Multiple comparison control was accomplished using the false discovery rate method of Benjamini and Hochberg with a q=0.2 (35). The experimental design is presented in **Supplementary Figure S1**. Human data was adjusted for medication and the differences in mean between the genotypes were assessed using ANCOVA, controlling for body mass index, post-mortem interval, age, education, sex, cognitive diagnosis, and beta-hydroxyisovaleroylcarnitine (selected as the measured acylcarnitines (AC) not influenced by APOE genotypes, to correct for general levels of acylcarnitines). The Tukey post-test was used to assess differences between genotypes. Significant variables were combined into cluster components as described above. ANCOVA analysis was performed on both individual variables and cluster components. Additionally, we used contrast post-test to investigate APOE genotype differences between individual metabolites in BCAA metabolic pathway.

## 3. Results

### 3.1 APOE genotypes influence the aging pattern of branch chain amino acids in mice brains

Metabolomic data show a high level of intercorrelation. To reveal the intercorrelation structure and to facilitate interpretation, data of each metabolite class were reduced using unsupervised variable clustering and converted into cluster components.

Variable clustering assigned 20 measured amino acids into 5 clusters. **Supplementary Table S2** contains a detailed cluster description, including the correlation between metabolites within each cluster. The age and APOE genotype-related differences in amino acid levels were not influenced by sex, which was tested using factorial models with sex x age and sex x genotype interactions. Therefore, further factorial analysis was performed with age, APOE genotype, and age x genotype interaction as main effects and sex as a covariate. The results from the factorial analysis are provided in **Supplementary Table S2**. Three clusters decreased with age: Cluster 1 containing alanine, asparagine, aspartic acid, glutamic acid, and threonine; Cluster 3 containing methionine, proline, and tyrosine; and Cluster 4 containing glycine, lysine, and serine (**Figure 1**). Clusters 3 and 4 showed an earlier decrease among APOE3/3 genotypes compared to the APOE2/2 and APOE4/4 genotypes, with the age x genotype interaction p=0.014 and p=0.09, respectively.

**Figure 1.**
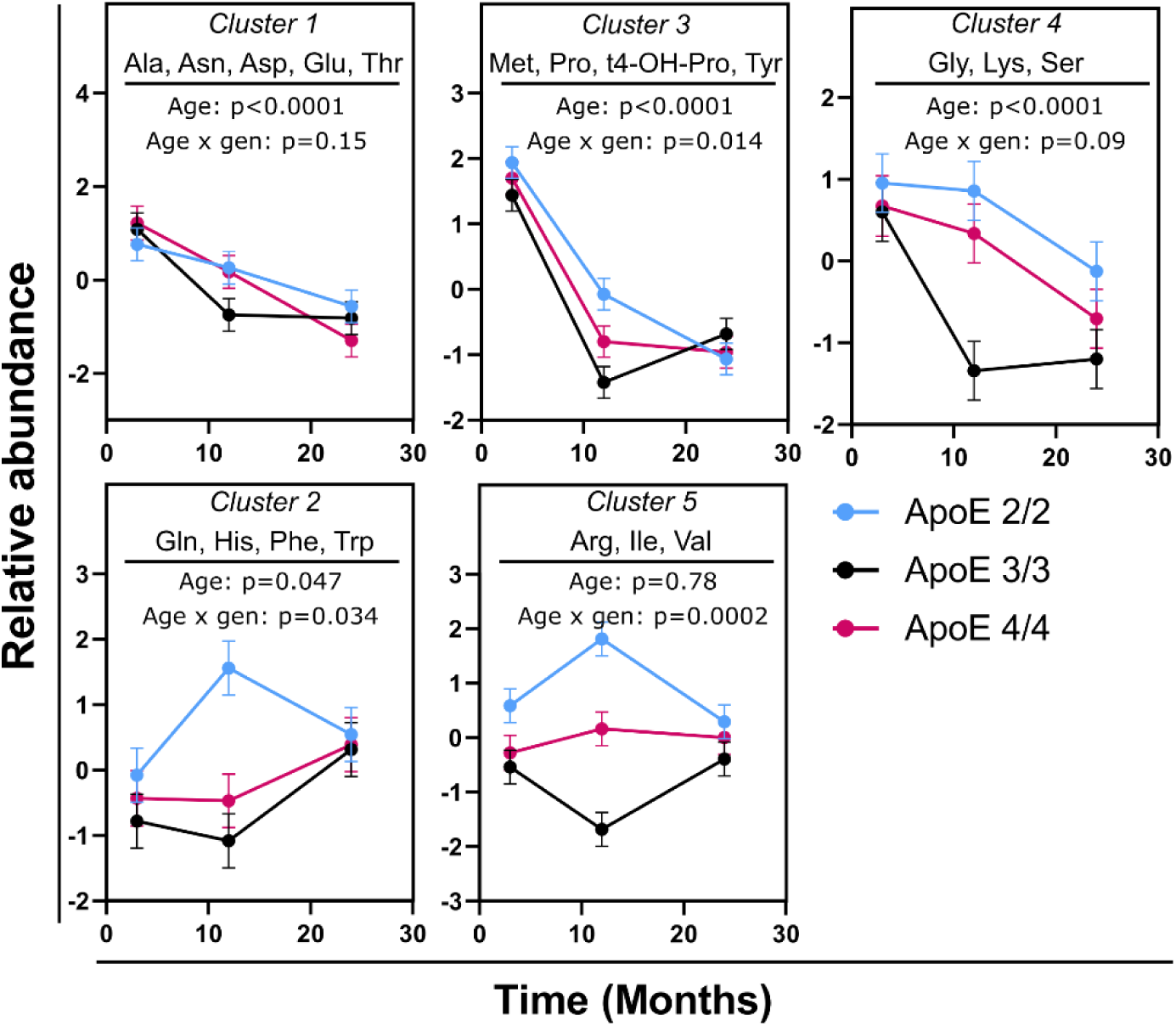
Differences in brain amino acids between mouse age groups, stratified by APOE genotype. Variables were converted into cluster components, with the members of each cluster indicated in each graph. Detailed description of each cluster is provided in **Supplementary Table S2**. Both males and females were used in the analysis. N=16 per genotype per time point (8 females and 8 males). Error bars represent standard errors. The indicated p-values were derived from a full factorial model testing for age, genotype, and age x genotype interaction. *Abbreviations:* Arg: Arginine; Ala: Alanine; Asn: Asparagine; Asp: Aspartic Acid; Gen: Genotype; Gln: Glutamine; Glu: Glutamic Acid; Gly: Glycine; His: Histadine; Ile: Isoleucine; Lys: Lysine; Met: Methionine; Phe: Phenylalanine; Pro: Proline; Ser: Serine; Thr: Threonine; Trp: Tryptophan; Tyr: Tyrosine; Val: Valine.

Cluster 2 containing glutamine, histidine, phenylalanine, and tryptophan, and Cluster 5 containing arginine, isoleucine, and valine, showed differential aging patterns among the genotypes (age x genotype, p_interaction_=0.034 and p_interaction_=0.0002, respectively). In both clusters, the APOE2/2 genotype showed an increase from 3 to 12 months (contrast post-test p=0.006 and p=0.006, respectively), followed by a decrease from 12 to 24 months (p=0.08 and p=0.0007, respectively). On the other hand, in Cluster 2, the APOE3/3 genotype showed no difference between 3 and 12 months and an increase from 12 to 24 months (p=0.02), and in Cluster 5, a decrease from 3 to 12 months (p=0.01) and then a restoration to the level of 3 months (p=0.004 for 12 vs. 24 months comparison). The APOE4/4 genotype showed no age-related changes in either cluster.

### 3.2. APOE genotypes influence the aging pattern of odd-chain AC in mice brains

Variable clustering assigned 35 measured AC and 29 calculated informative ratios and summations into 9 clusters, largely along their acyl chain length (**Supplementary Table S2**). Eight of those clusters showed age differences, and APOE genotype influenced the aging pattern of 5 clusters (significant age x genotype interaction) (**Figure 2**). Similar to the section above, no sex x age and sex x genotype interactions were detected, and sex was used as a covariate for this analysis.

**Figure 2.**
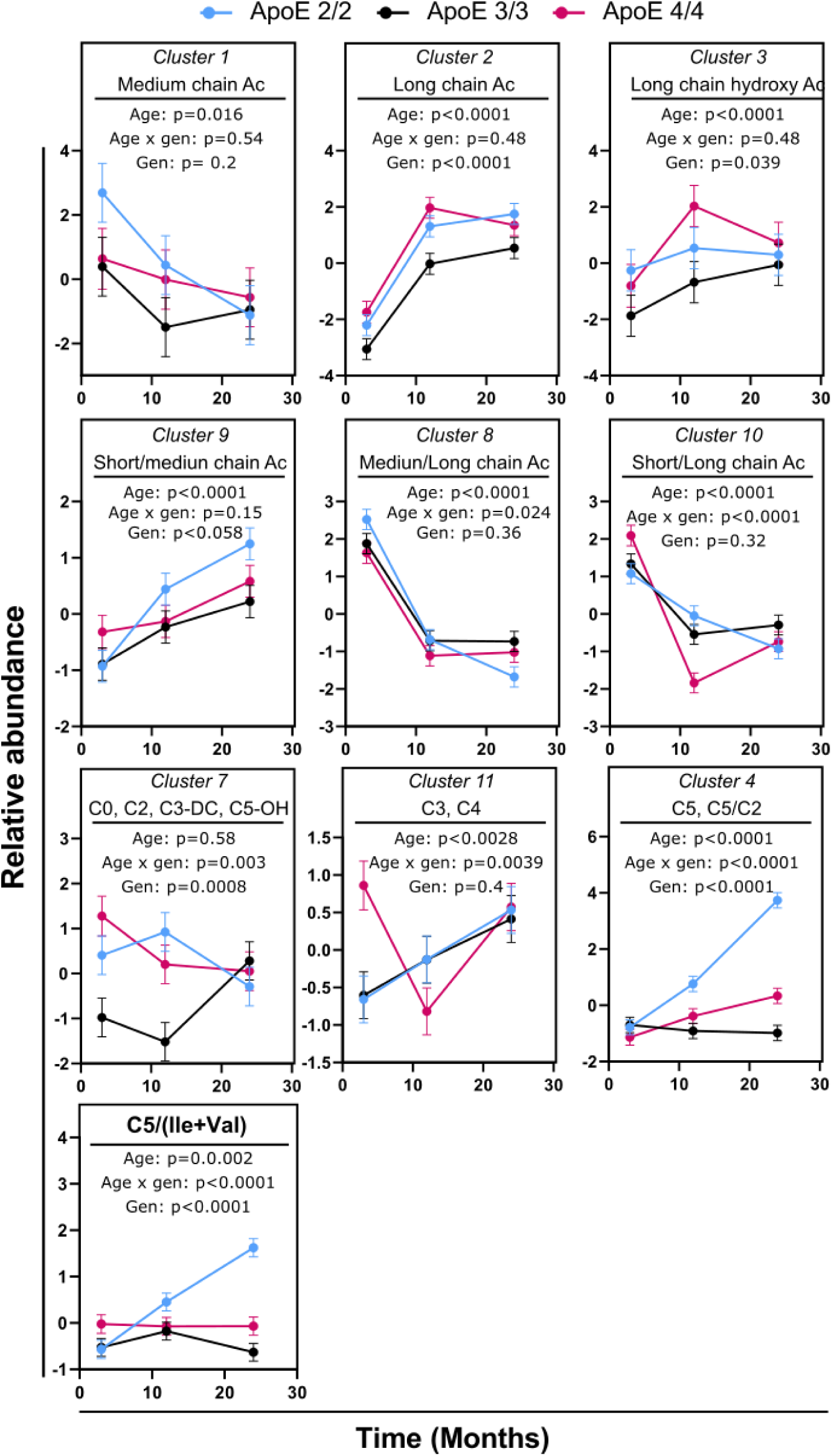
Differences in brain AC between mouse age groups, stratified by APOE genotype. Variables were converted into cluster components, with the members of each cluster indicated on each graph. Detailed description of each cluster is provided in **Supplementary Table S2**. Both males and females were used in the analysis. N=16 per genotype per time point (8 females and 8 males). Error bars represent standard errors. The indicated p-values were derived from a full factorial model testing for age, genotype, and age x genotype interaction. *Abbreviations:* AC: AC; Gen: Genotype; Ile: Isoleucine.

In general, the long-chain AC (LCAC, C12-C18, Clusters 2 and 3) increased with age, and medium-chain AC (MCAC, C6-C12, Cluster 1) decreased with age. Similarly, the ratio of MCAC to LCAC (Cluster 8) decreased with age. Additionally, LCAC were lower in the APOE3/3 genotype compared to the APOE2/2 and APOE4/4 genotypes; however, the ratio of MCAC to LCAC (Cluster 8) and short-chain to LCAC (Cluster 10) did not show differences between genotypes.

APOE genotype influenced the aging pattern of short-chain AC (Clusters 7, 11, and 4). In Cluster 7, represented by C0 and C2, the APOE3/3 genotype was higher at 24 months when compared to 3 and 12 months (p=0.004), with the other two genotypes showing no age-related differences. In Cluster 11, containing C3 and C4, the APOE3/3 and APOE2/2 genotypes were higher at 24 months when compared to 3 months (p=0.0006). On the other hand, the APOE4/4 genotype showed U-shaped age changes (3 vs. 12 months, p=0.0003; 12 vs. 24 months, p=0.002).

Cluster 4, containing C5 and the C5/C2 ratio, did not differ between genotypes at 3 months. It increased with age in the APOE2/2 genotype with a significant difference between 3 and 12 months (p<0.0001) and between 12 and 24 months (p<0.0001). The APOE4/4 genotype also showed an increase with age, with levels at 12 and 24 months higher than at 3 months (p=0.0014); however, this increase was of a lesser magnitude than the one observed in the APOE2/2 genotype, with APOE2/2 showing higher levels than APOE4/4 at 12 and 24 months (p=0.003 and p<0.0001, respectively). The APOE3/3 genotype did not show differences between the age groups.

C5 AC is derived from BCAA isoleucine (Ile), Leucine (Leu) and Valine (Val). To further investigate this part of metabolism, we calculated a product-to-substrate ratio (C5/(Ile+Val) as an insight into the performance of the metabolic pathway (**Figure 2**). The current Biocrates platform cannot distinguish between different C5 isoforms (i.e. valerylcarnitine, isovalerylcarnitine or 2-methylbutyrylcarnitine), products of different BCAA. Therefore, the sum of detected BCAA (Leu was not detected) was used in the calculation. The three genotypes showed no difference at 3 and 12 months. However, differential changes from 12 to 24 months were observed, with APOE2/2 showing an increase (p<0.0001), APOE3/3 showing a decrease (p=0.007), and APOE4/4 showing no change.

### 3.3 Biogenic amines show a uniform aging pattern among APOE genotypes

Variable clustering assigned 8 of the measured biogenic amines into 3 clusters (**Supplementary Table S2**). Two of those clusters (Cluster 1, containing the polyamines spermidine, spermine and putrescine; Cluster 3, containing symmetric dimethylarginine (SDMA) and carnosine) increased with age, and one cluster (Cluster 2, containing taurine, creatinine, and serotonin) decreased with age (**Figure 3**). Additionally, APOE genotype did not significantly influence the aging pattern of the clusters (no significant age x genotype interaction); however, significant differences between genotypes were detected at the 12-month time point for cluster 1 (APOE2/2 vs. APOE 4/4, p=0.001; APOE3/3 vs. APOE 4/4, p=0.003) and at the 24-month time point for cluster 2 (APOE2/2 vs. APOE 4/4, p=0.04; APOE2/2 vs. APOE 3/3, p=0.001). Similar to the section above, no sex x age and sex x genotype interactions were detected, and sex was used as a covariate for this analysis.

**Figure 3.**
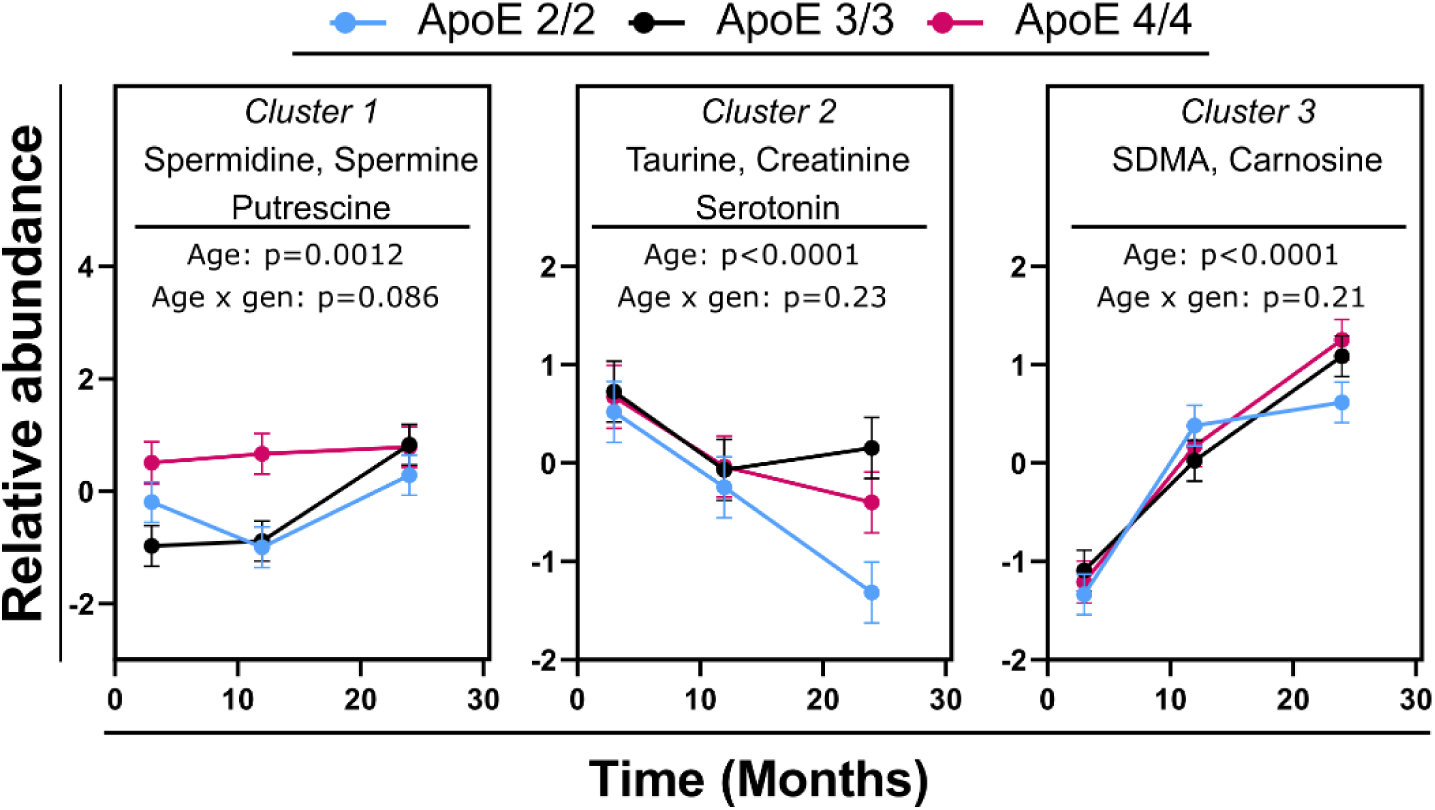
Differences in brain biogenic amines between mouse age groups, stratified by APOE genotype. Variables were converted into cluster components, with the members of each cluster indicated on each graph. Detailed description of each cluster is provided in **Supplementary Table S2**. Both males and females were used in the analysis. N=16 per genotype per time point (8 females and 8 males). Error bars represent standard errors. The indicated p-values were derived from a full factorial model testing for age, genotype, and age x genotype interaction.

### 3.4 Phosphatidylcholines show a uniform aging pattern among APOE genotypes

To better understand the changes in phosphatidylcholines (PC) in the aging brain, the age-related changes in 83 measured PC species were presented as a function of the sum of fatty acyl chains length and the number of unsaturated double bonds (**Figure 4**). For this analysis, we excluded a small number of species that showed genotype x age interaction (n=6) and the model was adjusted by sex and the APOE genotype. In general, the difference between the 3-month and 12-month time points was characterized by an increase in species with sum of carbons (C36–C40) and low unsaturated (1–3 double bonds). In contrast, there was a decrease in species that are highly unsaturated (≥4 double bonds) or have sum≥C42 or sum≤C32. The difference between the 12-month and 24-month time points was characterized by a further increase in low unsaturated species (1–2 double bonds), including species with a sum of C42 acyl chains, which decreased from the 3-month to the 12-month time point. This U-shaped age change is also shown in the cluster analysis (**Supplementary Figure S2**).

**Figure 4.**
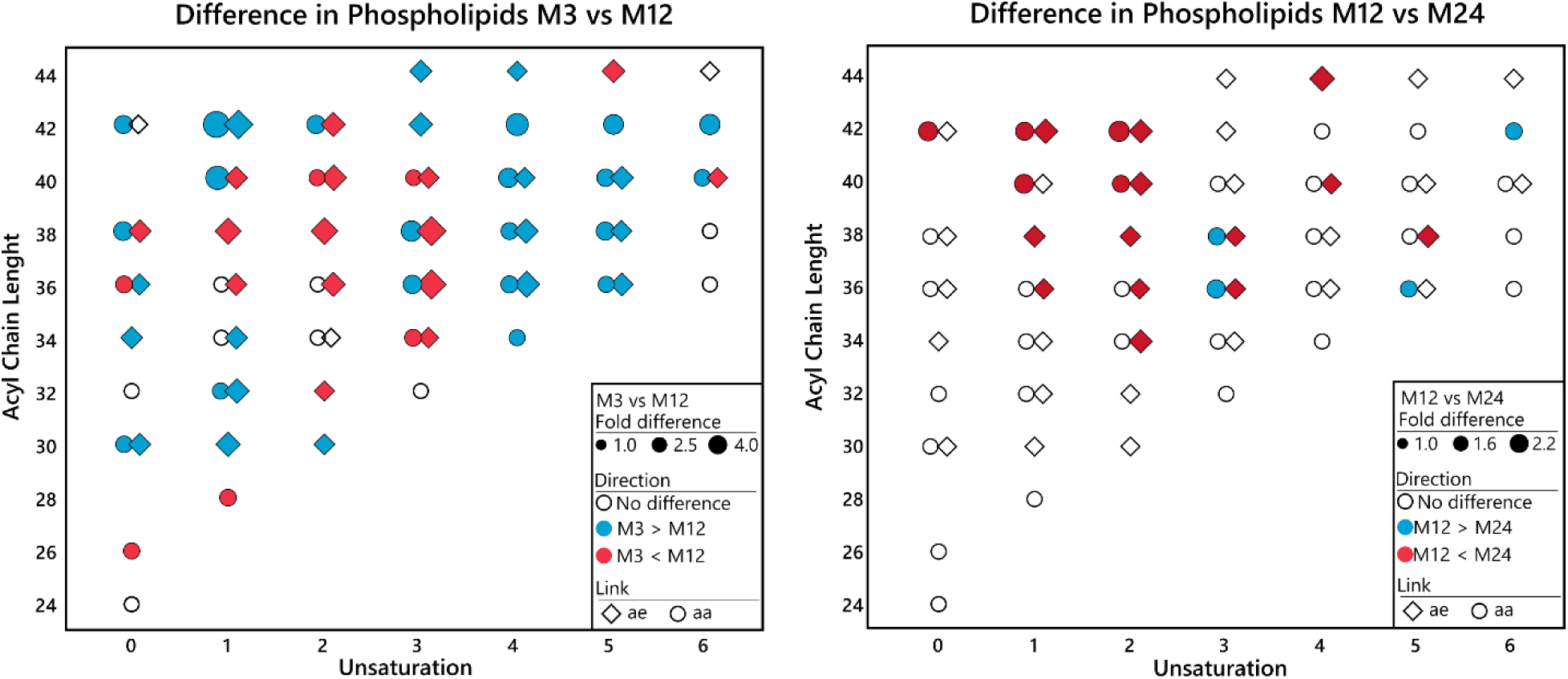
Differences in brain phosphatidylcholines between mouse age groups, emphasizing acyl chain length and the number of unsaturated double bonds. Differences with the p<0.05 and false discovery rate q=0.2 are marked with colors. Both males and females are used in the analysis. N=48 per time point.

### 3.5. Sphingomyelins show a sex-specific aging pattern

To facilitate data interpretation, SMs were clustered using the semi-unsupervised approach, with clustering performed separately for species that manifest similar age-related trends. This approach was applied since highly correlated metabolites were showing opposite aging trends. Out of 14 measured species, 10 species showed age-related differences and were converted into two clusters (**Figure 5 and Supplementary Table S2**). The Biocrates SM annotations were aligned with more in-depth SM species identification by targeted complex lipid platform generated by the Baker institute (both Baker and Biocrates annotations are present in the **Figure 5**) (10, 36). Baker SM annotations are used in the below results. Cluster 1, containing species with C18:0, C18:1 and C20:0 acyl chains decreased with age, and cluster 2 containing species with C16:0, C24:1 and 23:0 acyl chains increased with age. In addition, 5 species showed age x sex interactions, which warranted sex stratification of the analysis.

**Figure 5.**
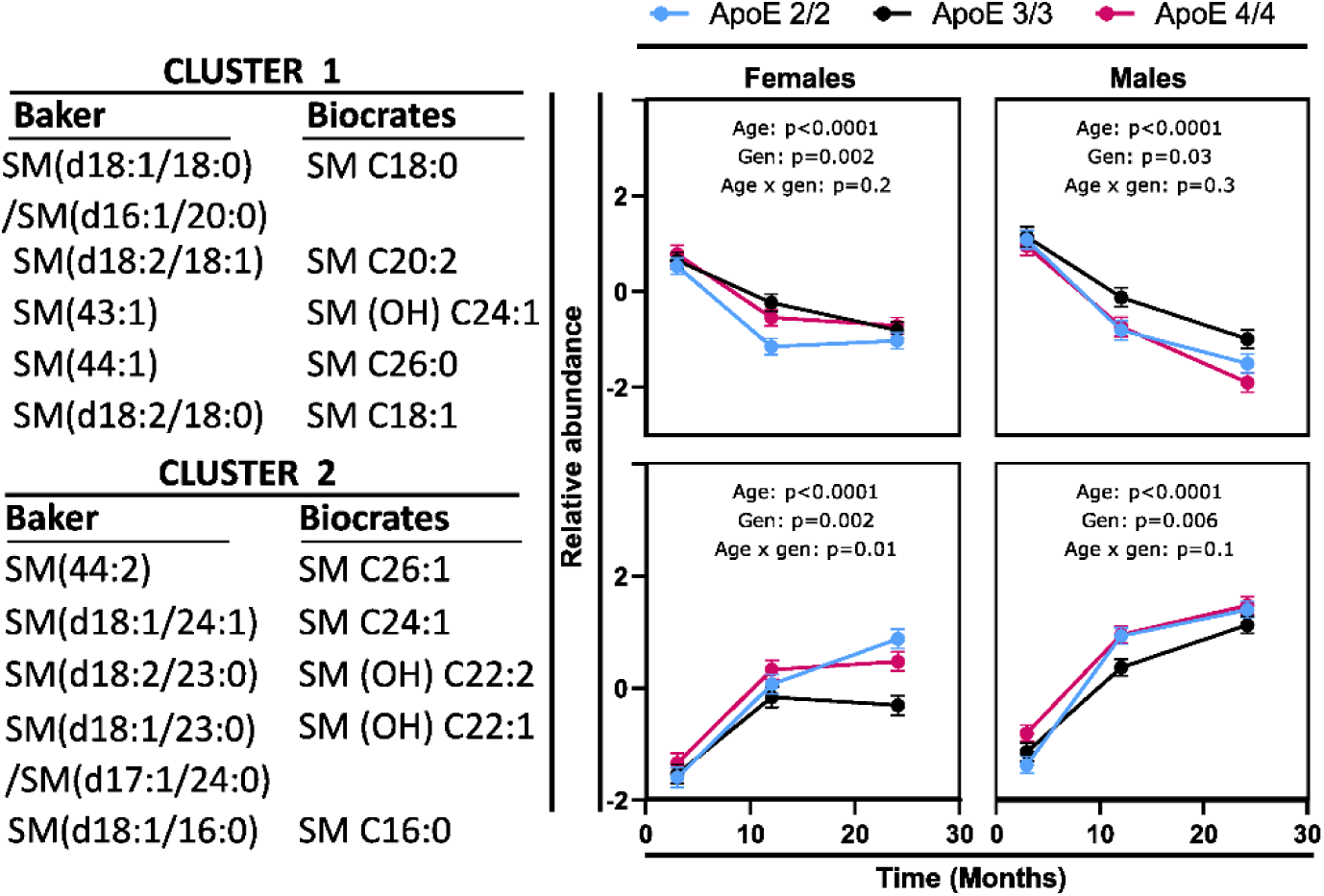
Differences in brain sphingomyelins between mouse age groups, stratified by sex and APOE genotype. Variables were converted into cluster components, with the members of each cluster indicated on each graph. Detailed description of each cluster is provided in **Supplementary Table S2**. N=8 per genotype, sex, and time point. The indicated p-values were derived from a full factorial model testing for age, genotype, and age x genotype interaction. Error bars represent standard errors. The Biocrates SM annotations were aligned with more in-depth SM species identification by targeted complex lipid platform generated by the Baker institute (10, 36).

In general, in females, a decrease in cluster 1 and an increase in cluster 2 occurred between 3 and 12 months, with no further differences between 12 and 24 months (post-test on the age effect 3 vs. 12 months, p<0.0001 for both clusters; 12 vs, 24 months, p=0.07 and p=0.1, respectively). On the other hand, age-related differences in males were observed between 3 and 12 months and between 12 and 24 months (3 vs. 12 months, p<0.0001 for both clusters; 12 vs. 24 months, p<0.0001 for both clusters). Neither cluster 1 nor cluster 2 showed age x genotype interaction.

### 3.6. Relevance to human metabolism

Next, we used previously published (16) human brain data from the ROS-MAP studies, generated using the Metabolon platform, to investigate similarities between the current mice model and human brain metabolism in the context of APOE genotype (ANCOVA results for all metabolites are presented in the **Supplemental Table S3**). For clarity of presentation, variable clustering was applied (**Figure 6A and Supplemental Table S4**). ROS-MAP brain metabolomic data were analyzed among the three most abundant APOE genotypes (APOE2/3, n=66; APOE3/3, n=304; and APOE3/4, n=112). Overall, we noticed the impact of APOE genotype on potential products of β-oxidation and relevant BCAA metabolism. Compared to APOE3/3 and/or APOE3/4 carriers, APOE2/3 carriers had a lower level of carnitine and AC, including (S)-3-hydroxybutyrylcarnitine, eicosenoylcarnitine (C20:1), arachidonoylcarnitine (C20:4), acetylcarnitine (C2), and tiglyl carnitine (C5:1) (an intermediate in BCAA metabolism (37)). In addition, compared to APOE3/3, APOE2/3 carriers had lower levels of a cluster composed of several markers of impaired BCAA utilization for energy, including methylmalonate and 3-methylglutaconate (inversely correlated with BCAA utilization in the TCA cycle (38)) and alpha-hydroxyisovalerate produced by incomplete catabolism of BCAA (39)).

**Figure 6.**
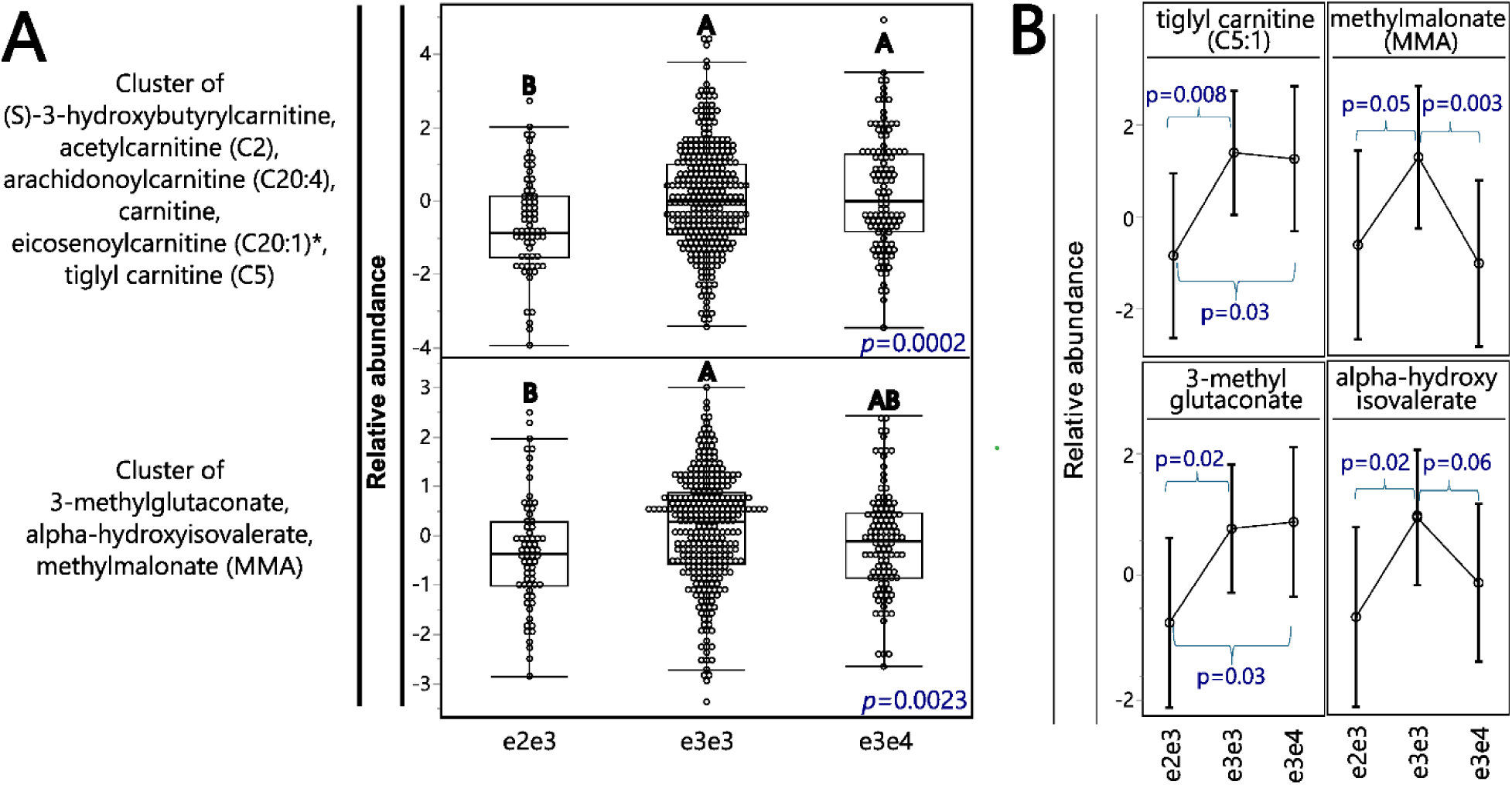
APOE genotypes influence brain AC and branch-chain amino acids (BCAA) metabolism in the Religious Orders Study/Memory and Aging Project (ROS-MAP) cohort. **A)** Differences between cluster components among APOE genotypes. Cluster components were generated using metabolites manifesting APOE genotype differences and analyzed with ANCOVA followed by the Tukey pos-hoc test, with the significance indicated by letters. N: (e2e3= 66; e3e3= 304; e3e4= 112). **B)** APOE genotype differences among individual metabolites involved in BCAA metabolism. Error bars represent standard error of the mean. P values are derived from the contrast post-test.

There are several metabolic pathways that can bring carbon from BCAA to the TCA cycle (pathways reviewed and illustrated in (37, 40)). Therefore, in addition to cluster analysis, we have analyzed the APOE genotype differences in levels of individual markers of BCAA metabolism (**Figure 6B**). In general, we observed two patterns of differential metabolite levels among APOE genotypes. First, the tiglyl carnitine and 3-methylglutaconate were lower in APOE2/3 carriers, compared to both APOE3/3 and APOE3/4 (p_values_ as follow for tiglyl carnitine and 3-methylglutaconate: 2/3 vs 33 =0.008 and 0.02; 2/3 vs 3/4 =0.03 and 0.03. The p_value_ was derived using contrast post-test). Second, the alpha-hydroxyisovalerate and methylmalonate were lower in both APOE2/3 and APOE3/4 carriers, when compared to APOE3/3 (p_values_ as follow for alpha-hydroxyisovalerate and methylmalonate: 2/3 vs 33 =0.02 and 0.05; 3/4 vs 3/3 =0.06 and 0.003. The p_value_ was derived using contrast post-test).

In contrast to the rodent model, BCAA level remained the same among all human genotypes (data not shown). Furthermore, APOE2/3 carriers had lower levels of kynurenate and kynurenine, part of the tryptophan metabolism via the kynurenine pathway (**Supplementary Table S3, Figure 7**), however tryptophan did not differ among genotypes (data not shown). Relevantly, N-acetylglucosamine/N-acetylgalactosamine was higher in APOE2/3 carriers compared to APOE3/4 carriers; it also showed a higher trend compared to APOE3/3 carriers, though it was not significant.

**Figure 7.**
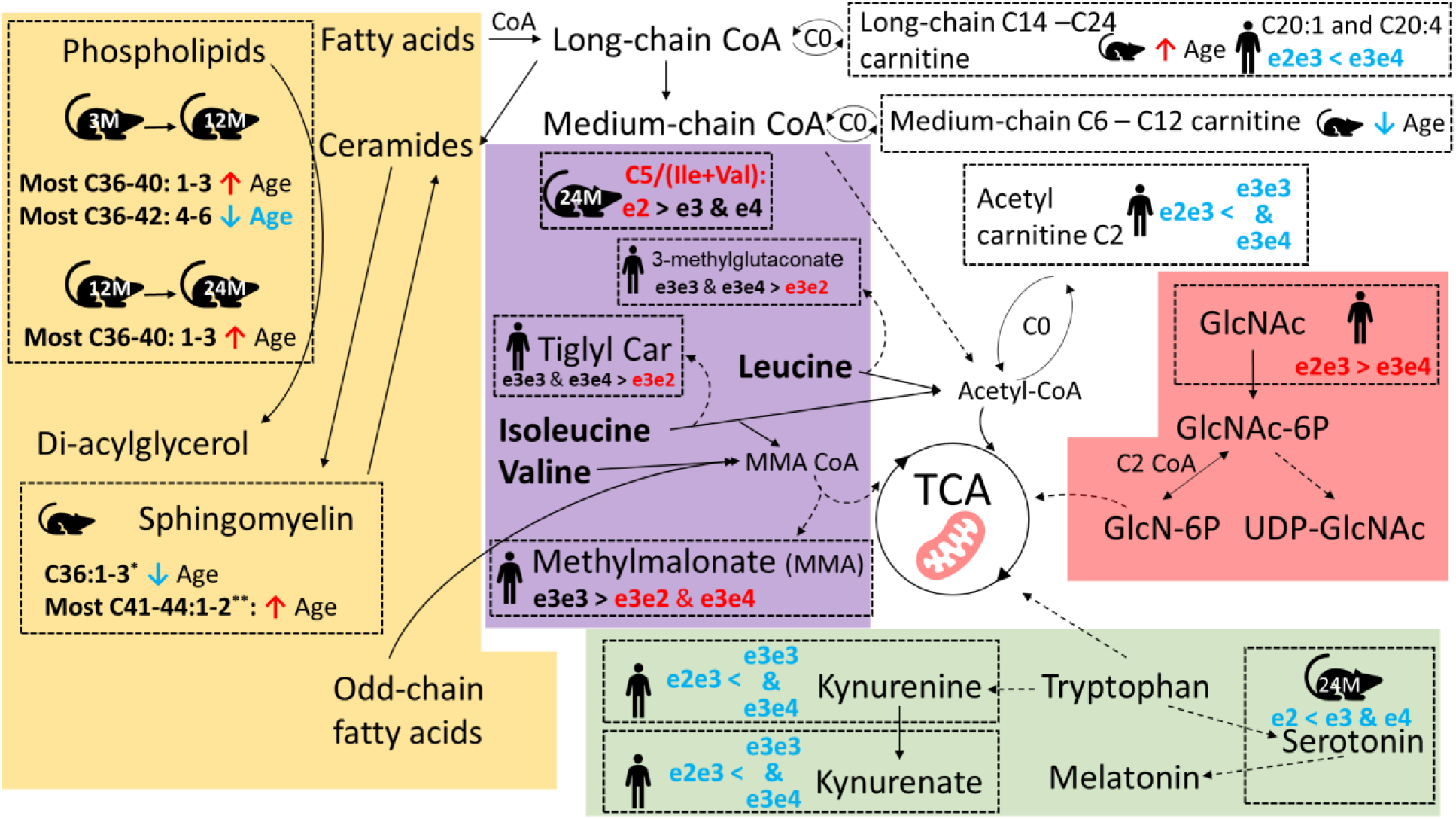
APOE genotypes- or age-related brain energy metabolism pathways highlighting different substrates for the tricarboxylic acid (TCA) cycle, with the summary of findings from mice and human data utilized in the current manuscript. *: C36:1-3 sphingomyelin includes SM(d18:1/18:0)/SM(d16:1/20:0), SM(d18:2/18:0) and SM (d18:2/18:1). **: Most C41-44:1-2 sphingomyelin includes SM (d18:1/23:0)/SM(d17:1/24:0), SM (d18:2/23:0), SM (d18:2/24:1), and SM (44:2).

## 4. Discussion

In the current manuscript, we conducted targeted metabolomics to characterize the aging brain metabolism of the mice model that carries human APOE2/2, APOE3/3, and APOE4/4 genes. Using Biocrates P180 platform, we have provided absolute quantifications of ∼ 150 metabolites and informative rations, broadly covering phospholipids, sphingolipids, acylcarnitines, biogenic amines and amino acids. Our analysis identified that the AD protective APOE2/2 genotype manifests elevated markers of branch chain amino acid utilization in β-oxidation in mice brain. Additionally, we showed evidence that the APOE2/3 and APOE3/4 genotypes influence β-oxidation and BCAA in human brain, although the phenotypic manifestations of this involvement were different. Moreover, we described lipidomic and metabolomic aging patterns of the current mice model, with several key metabolites serving as aging markers in humans, also linked to AD. Together, these results suggest a potential involvement of the APOE2 genotype in energy metabolism among others and characterize the current mice model for future study of APOE in AD, brain aging, and brain BCAA utilization for energy. The observed age and APOE genotype differences and their connection to energy metabolism are summarized in **Figure 7**.

APOE4 is associated with hypometabolism of glucose in the brain, possibly occurring well before disease onset (41). Together with other metabolic alterations, such as accumulation of lactate observed in human AD brains (42–45), and decreased transport of pyruvate to the mitochondria, low TCA cycle productivity is expected (26), causing mitochondrial dysfunction and energy deficits. Such circumstances trigger compensatory metabolism seeking alternative energy sources. Such a phenomenon may be reflected in various metabolites that can contribute substrates to the TCA cycle, such as amino acids, keto-acid derivatives, ketone bodies, fatty acids, and acyl carnitines. TCA cycle deficits can also affect phospholipids synthesis and other anabolic processes relying on its functioning.

In the current mice model, only those with the APOE2/2 genotype had brain levels of BCAA and tryptophan showing a parabolic association with age, potentially suggesting an increase in BCAA and tryptophan utilization in older age. This contrasts with the gradually increasing level of these amino acids with age in the brains of APOE3/3 and APOE4/4 mice. BCAA can be oxidized into BC-keto acids, further transformed into BC-acylCoAs, then to acetylCoA that can enter the TCA cycle. Interestingly, the first conversion step can be coupled with alpha-ketoglutarate (from TCA cycle) and produce glutamate (46). Such a reaction doesn’t only consume TCA cycle intermediates but may also interfere with the balance of the excitotoxic glutamate. In addition, BCAA compete on transfer through the BBB with other large neutral amino acids, namely aromatic amino acids (Trp, Phe, Tyr). BCAA are involved in synthesis of neurotransmitters, proteins and energy production, therefore their effects reach farther than the described here and have greater implication in AD (47). Lower blood levels of BCAA were a main contributor to predicted risk of dementia and AD in several cohorts, even 10 years before disease onset (48–51). On the contrary, high brain BCAA levels were associated with AD pathology and cognitive impairment (27), suggesting the importance of their utilization by brain. For tryptophan, the above results partially align with human findings demonstrating decreased blood levels in people with AD (52–54), and with aging (55, 56). In a small study, tryptophan was increased in five brain regions in people with AD compared to control (n=9 only, (43)).

Similarly, mice with the APOE2/2 genotype showed an increase in the ratio C5 AC/(Ile+ Val) (product to indirect precursor), further supporting the potential increase in utilization of BCAA in TCA. In comparison, mice with APOE4/4 genotype had milder increases in short-chain acyl carnitines at the age of 12 months, suggesting that protein catabolism and oxidation of amino acids (or fatty acids) was less pronounced, hypothetically less successful in compensating the already reduced energy production.

Analysis of human brain DLPFC data has consistently shown that APOE2/3 genotype has lower levels of the above markers of alternative energy production. These included acetylcarnitine (C2), the terminal product of beta oxidation and the main shuttle of fatty acids (can cross the BBB and has other functions/effects (57)). Importantly, APOE2/3 genotype also had lower levels of several markers and intermediates in BCAA metabolism (pathways of BCAA metabolism leading to the TCA cycle are reviewed in (37, 40)). Those included: the tiglyl carnitine, an intermediate in BCAA metabolism (37), which plasma and urine levels positively associated with inborn error of isoleucine metabolism and resulting neurodegeneration (58); 3-methylglutaconate, inversely correlated with BCAA utilization in the TCA cycle (38) with elevated urine levels positively associated with inborn error of leucine metabolism and resulting neurodegeneration (59). On the other hand, methylmalonate and alpha-hydroxyisovalerate were lower in both APOE2/3 and APOE3/4, compared to APOE3/3. Methylmalonate CoA is one of the final steps in valine and some branch chain fatty acids metabolism, and inborn errors in this pathway result in elevated methylmalonate levels (38). Alpha-hydroxyisovalerate was inversely associated with dietary BCAA intake (39). BCAA can enter TCA cycle along three enzymatic pathways, with methylmalonate-CoA or acetyl-CoA as the final products, ready to be incorporated into the TCA (37, 40).

Leucine enters TCA cycle through acetyl-CoA, valine through methylmalonate CoA and isoleucine through both. The above results could suggest that in human brain, APOE2 allele elevates metabolism of all three BCAA (and potentially branch chain fatty acids), whereas APOE4 allele elevates metabolism of only valine (or branch chain fatty acids). As those results do not explain the derogatory effect of APOE4 allele, they can suggest how APOE2 allele can be protective against neurodegeneration.

Human APOE2/3 carriers also had lower brain levels of tryptophan metabolites (i.e., kynurenate and kynurenine in DLPFC), a pathway that is implicated in cognitive decline (60). Kynurenine can cross the BBB while kynurenic acid cannot, and both can be produced in the brain from Trp (61). Like other metabolites in the kynurenine pathway, their neuroactivity mainly stems from modulation of inflammation via activation of the aryl hydrocarbon receptor (AhR) (62). While evidence of link to AD in human CSF and blood are inconsistent, kynurenine was elevated in the frontal cortex of people who died with AD, compared to controls (42). Another metabolic perturbation observed in human APOE2/3 carriers included higher levels of N-acetylglucosamine or N-acetylgalactosamine (those metabolites cannot be distinguished with the current Biocrates platform) in DLPFC, a human or microbial precursor of uridine-diphosphate-N-acetylglucosamine (UDP-GlcNAc), which also modulates the uptake of acetyl-CoA via the hexosamine biosynthetic pathway (63, 64). However, DLPFC brain levels of N-Acetyl-glucosamine 1-phosphate (UDP-GlcNAc direct precursor) are strongly associated with cognitive decline and Tau tangles in the same cohort (26). The above findings in both rodents and humans suggested that APOE2 increases the level of brain energy metabolism compared to other genotypes in older ages, while also highlighting differentially regulated processes.

Additionally, in the current manuscript, we report that the metabolic signature of aging mice brain with alternation of these aging signatures is linked to susceptibility/resistance to cognitive decline in humans. For example, our findings indicate that in the current mice model, long-chain phospholipids (C36-42) and highly unsaturated phospholipids (4-6 double bonds) tend to decrease with age. Consistently, in human brain DLPFC, the highly unsaturated long-chain phospholipids – such as 1-linoleoyl-2-arachidonoyl-GPC (18:2/20:4n6) (PC C38:6) and 1-oleoyl-2-docosahexaenoyl-GPC (18:1/22:6) (PC C40:7) – showed a strong negative association with cognitive decline (15). Therefore, data in both mice and human support that a higher level of these phospholipids persisting in brains is associated with resilience to metabolic aging. This could be related to the reduced TCA productivity affecting phospholipid synthesis, resulting in low neurogenesis and decreased synaptic plasticity.

It is well established that oxidative stress and inflammation are part of the deleterious processes involved in neurodegeneration. Oxidative stress affects the TCA cycle and can be linked to increased brain fructose metabolism (65, 66), which is affected by impaired glucose metabolism in APOE4 carriers (67, 68). Moreover, mouse model of AD showed higher vulnerability to amyloid-beta induced oxidative stress in carriers of APOE4 isoform, since they lack strategic Cys residues (replaced by Arg) in the APOE protein, which can attack free radicals and limits lipid peroxidation (69, 70). Thus, it increases the rate of neuronal modification, damage, and death. An environment of oxidative stress and inflammation increases post-translational modification of proteins and accelerates proteolysis and release of modified amino acid residues. For example, SDMA is formed via methylation of the arginine residue of a protein, and as such could increase with oxidative stress and cellular damage. We observed that brain SDMA is positively associated with age in our mouse model, and SDMA content in human brain DLPFC was also positively associated with cognitive decline (15). SDMA also increased in frontal cortex brain tissue of people who died with AD (42). Therefore, the additional accumulation of SDMA in brain may indicate a high level of susceptibility to metabolic aging in addition to the process of normal aging, which may have implications in neurogenerative diseases at advanced ages. Indeed, SDMA was associated with an array of health conditions, including muscle mass loss (71), renal failure (72) and neurodegeneration (73) including vascular dementia and AD (74). Enzymatic arginine metabolism is also altered in AD brains, showing decreased activity and protein expression of NO synthase and increased arginase (74, 75). We found that arginine metabolism into polyamines (putrescine, spermidine, and spermine) increased with age in mice brain (with earlier increase in APOE4 carriers). While depending on measured brain region, putrescine was decreased in the brain of people with AD (75, 76), and mice model of AD (77), while spermine and spermidine increased in human AD brains (76, 78, 79). Although polyamines act as reactive oxygen species (ROS) scavengers, they affect age-related conditions and can be toxic (80). In excess, spermidine and spermine may cause NMDA receptor excitotoxicity. Also, they can be further metabolized to aldehydes and then to the toxic acrolein, which binds to proteins and nucleic acids with detrimental effects (80, 81). Moreover, amyloid beta may cause an increase in polyamine metabolism manifested by up-regulated spermine uptake and increased ornithine decarboxylase that converts ornithine to putrescine (82, 83). Other amines also act as antioxidants, for example taurine (decreased in our mice model with age and with APOE2 genotype), an amino acid that can be endogenously produced from cysteine using vitamin B6. It acts as a non-excitatory neurotransmitter and may be neuroprotective (84). Another example is carnosine (decreased in our mice model with age and with APOE2 genotype), a dipeptide (beta-alanine - histidine) hence could on one hand indicate post-translational modification, but on the other hand is an antioxidant and provides intracellular pH buffering (85). The above findings have highlighted the similarities of the human brain metabolome and the current mice model brain metabolome, which both point to the potential pathways involved in the resilience/susceptibility to metabolic aging, a complicated process implicated in neurodegenerative diseases (86).

In contrast, we have also noticed the differences between the mice model and the human brain age-related metabolome. For example, while biogenic amines like the neurotransmitter serotonin decreased with age in the current mice model and other rodent models (87, 88), the serotonin levels remained the same in aging human brain (89). Nevertheless, the serotonin/Trp ratio (marker for serotonin synthesis) decreased in frontal cortex brain tissue of people with AD, compared to control (42). In the aging human brain, serotonin metabolism is impaired by the altered activity levels of serotonin transporter and receptor, both are further implicated in neurodegenerative diseases (90–92).

Complex lipids also showed significant changes in our study. PC and SM in the mice model points to the exchange of fatty acids between these two pools via the ceramide pathway: SM C36:1 is a major SM species in mice brain (93) and its high concentration is highly specific to mice brain compared to other tissues (94). In the current mice model, SM C36:1 decreased with age and PC C36:1 increased with age. As SM is a major component in neuronal myelin (94–96), our observation of decreasing dominant SM species with age is consistent with the “myelin hypothesis” of normal and abnormal aging, in which the production and maintenance of myelin are compromised over time during these processes (97, 98). The accompanying opposite trend of PC C36:1 can reflect the reduced production of SM via the ceramide pathway. In contrast, in aging human brains, PC and SM – except for a few differences in PC – remain largely unchanged with aging. These differences between human and mice data can be due to the species-specific metabolism, or the fact that human brain samples are limited to a narrower sample collection time frame and to the brain regions available (99). Despite these differences, compared to available human brain metabolome data, our findings in the mice model point to shared pathways of mice and human brain metabolome in terms of normal and abnormal aging. This enables the further utilization of this mice model to investigate the nature of metabolic aging and the age-associated diseases in the human central nervous system.

In conclusion, findings here have documented the APOE and aging brain metabolic signatures of a mice model of AD, which points to the effect of APOE on brain energy metabolism via regulation of amino acid and acyl carnitine pathways. This model enables further studies on APOE’s effect on age-related neurodegenerative diseases of humans and on their potential treatments. It is also important to acknowledge that our understanding of brain energy metabolism in the context of those metabolic markers is very limited, and the current results are only speculative. However, this manuscript points towards a need for more in-depth analysis where the impact of APOE genotype is understood on the cellular level and can be further translated to the whole brain metabolic alteration.

## Limitations

The limited available human data allowed us to compare only the haplotype for APOE isoforms. Additionally, all results here describe associations with causality to be further confirmed. In addition, identified C5 acyl carnitine does not distinguish between specific isoforms, limiting data interpretation.

## Acknowledgements

The Alzheimer’s Disease Metabolomics Consortium (ADMC) is funded wholly or in part by the following National Institute on Aging grants and supplements, which are components of the Accelerating Medicines Partnership for AD (AMP-AD) and/or Molecular Mechanisms of the Vascular Etiology of AD (M2OVE-AD): NIA R01AG081322, R01AG046171, 1RF1AG051550, 1RF1AG057452, R01AG059093, 1RF1AG058942, U01AG061359, U19AG063744, 3U19AG063744-04S1, and 1R01AG081322 awarded to Dr. Rima Kaddurah-Daouk at Duke University in partnership with a large number of academic institutions. Dr. Guojun is supported by RF1AG051504. Dr. Arnold is supported by R01AG069901. We wish to acknowledge the editing assistance of Jon Kilner, MS, MA (Pittsburgh).

## Author Contributions

K.B. and N.L. analyzed and interpreted the data and wrote the manuscript. N.L. N.Z., T.K., and G.B. designed and executed the animal studies by generating experimental mice and harvesting serum for metabolomics analyses. N.K., KH contributed to manuscript writing and data interpretation. M.A., S.M., A.K. performed data preprocessing and contributed to data analysis. RK-D. is a principal investigator of this project and was engaged in the overall design and implementation of the project and secured the necessary funds.

## Data Availability Statement

All human data (ROS-MAP) used in this paper are available via the AD Knowledge Portal (https://adknowledgeportal.org) and also through the RADC Research Sharing Hub (https://www.radc.rush.edu/). The AD Knowledge Portal is a platform for accessing data, analyses, and tools generated by the Accelerating Medicines Partnership (AMP-AD) Target Discovery Program and other National Institute on Aging (NIA)-supported programs to enable open-science practices and accelerate translational learning. The data, analyses and tools are shared early in the research cycle without a publication embargo on secondary use. Data is available for general research use according to the following requirements for data access and data attribution (https://adknowledgeportal.org/DataAccess/Instructions). All generated mice data will be publicly available at: https://www.synapse.org/#!Synapse:syn20808172.

## Competing Interests

RK-D is an inventor on several patents in the field of metabolomics and holds equity in Metabolon Inc., Chymia LLC and Metabosensor, all of which were not involved in this study. M.A. is an inventor on several patents on applications of metabolomics in diseases of the central nervous system; he holds equity in Chymia LLC, which were not involved in this study. G.B. is a consultant for SciNeuro Pharmaceuticals and Kisbee Therapeutics. He is also the Editor-in-Chief of Molecular Neurodegeneration. All other authors declare no competing interests.

