## Supplementary figures and images for "APOE Genotype Influences on The Brain Metabolome of Aging Mice – Role for Mitochondrial Energetics in Mechanisms of Resilience in APOE2 Genotype"

### Supplementary Figure S1

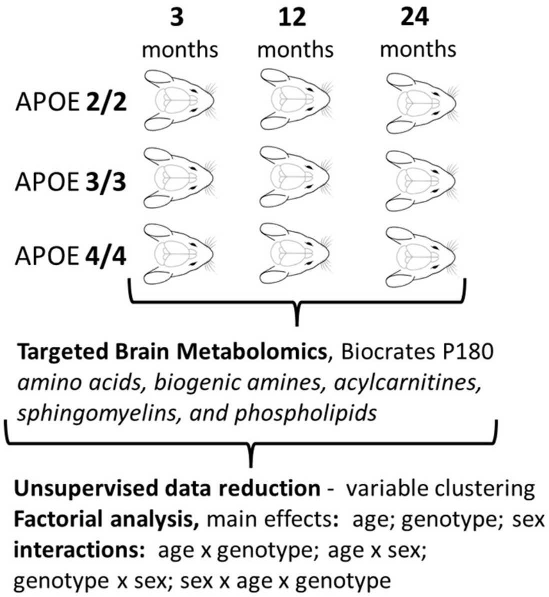


**Figure S1. Experimental design.**
