## Supplementary Figure S2 for "APOE Genotype Influences on The Brain Metabolome of Aging Mice – Role for Mitochondrial Energetics in Mechanisms of Resilience in APOE2 Genotype"

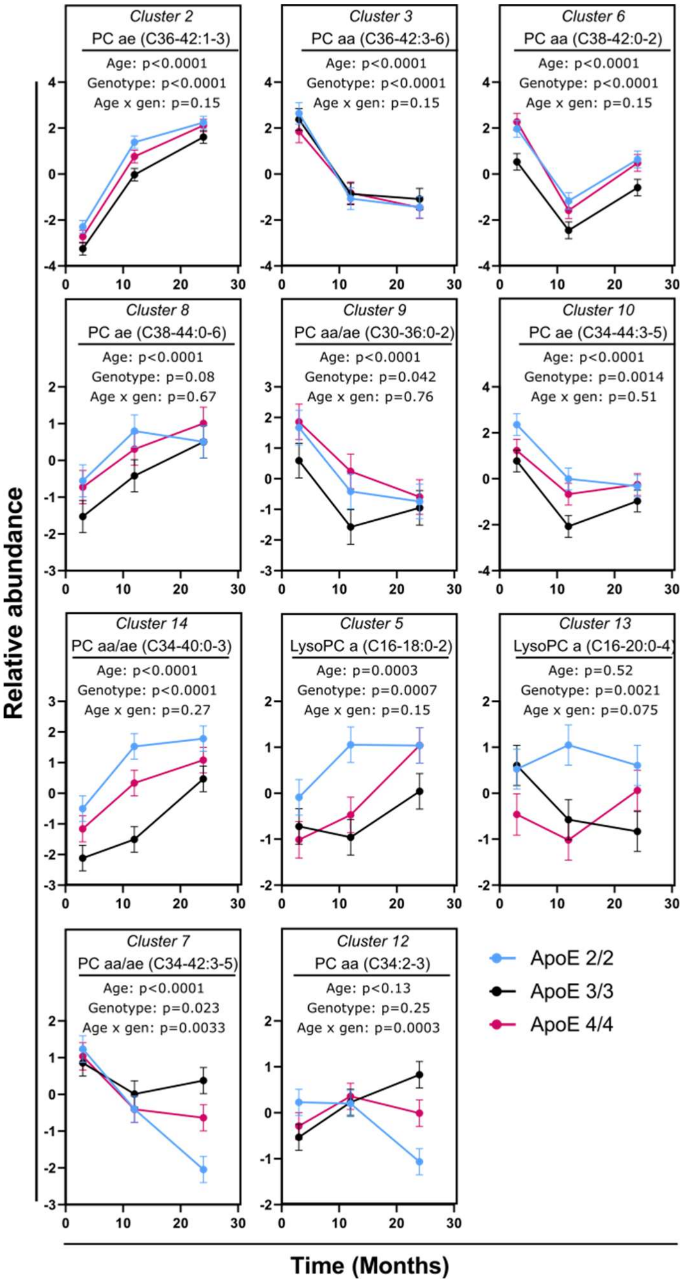


**Figure S2. Aging pattern of phospholipids between APOE genotypes.** To reveal the intercorrelation structure and to facilitate interpretation, data were reduced using unsupervised variable clustering and converted into cluster components. Supplementary Table S2 contains a detailed cluster description, including the correlation between metabolites within each cluster.
