## Supplementary Table S1 for "APOE Genotype Influences on The Brain Metabolome of Aging Mice – Role for Mitochondrial Energetics in Mechanisms of Resilience in APOE2 Genotype"

**Table S1. The demographics of the ROS-MAP cohort, subset used for the analysis.**

| | ApoE2/3 | ApoE3/3 | ApoE 3/4 |
|---|---|---|---|
| n= | 66 | 304 | 112 |
| Age at death (mean ± std eva) | 92.3±6.6 | 90.5±6.35 | 89.5±5.5 |
| BMI (lastest data available) | 24.1±5.08 | 26.2±5.28 | 25.4±4.54 |
| Education | 15.4±3.11 | 15.9±3.41 | 16.3±3.24 |
| PMI | 7.51±4.01 | 8.5±5.36 | 7.29±3.91 |
| Sex (Male %) | 18.18 % | 31.25 % | 32.14 % |
| Final consensus cognitive diagnosis | | | |
| NCI: No cognitive impairment (No impaired domains) | 41% | 32% | 18% |
| MCI: Mild cognitive impairment (One impaired domain) and NO other cause of CI | 17% | 27% | 19% |
| MCI: Mild cognitive impairment (One impaired domain) AND another cause of CI | 5% | 0% | 1% |
| AD: Alzheimer's dementia and NO other cause of CI (NINCDS PROB AD) | 30% | 35% | 52% |
| AD: Alzheimer's dementia AND another cause of CI (NINCDS POSS AD) | 8% | 5% | 8% |
| Other dementia: Other primary cause of dementia | 0% | 2% | 3% |
