## Supplementary Table S2 for "APOE Genotype Influences on The Brain Metabolome of Aging Mice – Role for Mitochondrial Energetics in Mechanisms of Resilience in APOE2 Genotype"

**Supplemental Table S2.** Describtion of cluster analysis with each cluster membership. P values from factorial analysis performed on individual metabolites and cluster components.

| Class | Compound | Cluster information | | | | P values - Full factorial model on individual metabolites | | | | | | | P values - Full factorial model on cluster components | | | |
|---|---|---|---|---|---|---|---|---|---|---|---|---|---|---|---|---|
| | | Cluster | RSquare with Own Cluster | RSquare with Next Closest | 1-RSquare Ratio | Age | Genotype | Gender | Age*Gender | Genotype*Age | Genotype*Age *Gender | Genotype*Gen der | Age | Gender | Genotype | Genotype*Age |
| Amino Acids | Asn | | 0.73 | 0.40 | 0.46 | 0.0001 | 0.53 | 0.17 | 0.74 | 0.0043 | 0.65 | 0.71 | | | | |
| | Glu | | 0.70 | 0.32 | 0.44 | 0.0001 | 0.22 | 0.39 | 0.87 | 0.92 | 0.3 | 0.13 | | | | |
| | Asp | **1** | 0.44 | 0.11 | 0.63 | 0.0029 | 0.042 | 0.2 | 0.034 | 0.012 | 0.88 | 0.67 | 0.0001 | 0.11 | 0.55 | 0.15 |
| | Thr | | 0.41 | 0.22 | 0.75 | 0.0001 | 0.07 | 0.0019 | 0.56 | 0.092 | 0.86 | 0.57 | | | | |
| | Ala | | 0.35 | 0.27 | 0.90 | 0.45 | 0.13 | 0.52 | 0.37 | 0.021 | 0.71 | 0.98 | | | | |
| | Phe | | 0.86 | 0.53 | 0.29 | 0.6 | 0.0001 | 0.053 | 0.53 | 0.0005 | 0.54 | 0.35 | | | | |
| | His | **2** | 0.83 | 0.27 | 0.23 | 0.82 | 0.046 | 0.22 | 0.27 | 0.02 | 0.16 | 0.067 | 0.047 | 0.83 | 0.0018 | 0.034 |
| | Gln | | 0.79 | 0.18 | 0.25 | 0.0001 | 0.098 | 0.92 | 0.38 | 0.6 | 0.48 | 0.049 | | | | |
| | Trp | | 0.64 | 0.18 | 0.44 | 0.0001 | 0.0027 | 0.0053 | 0.6 | 0.032 | 0.66 | 0.29 | | | | |
| | Met | | 0.79 | 0.28 | 0.29 | 0.0001 | 0.0001 | 0.25 | 0.11 | 0.5 | 0.096 | 0.3 | | | | |
| | Tyr | **3** | 0.76 | 0.22 | 0.31 | 0.0001 | 0.16 | 0.0001 | 0.058 | 0.11 | 0.1 | 0.26 | 0.0001 | 0.0001 | 0.048 | 0.014 |
| | Pro | | 0.69 | 0.47 | 0.60 | 0.0001 | 0.025 | 0.0035 | 0.59 | 0.0001 | 0.24 | 0.46 | | | | |
| | t4-OH-Pro | | 0.37 | 0.18 | 0.77 | 0.0001 | 0.0004 | 0.0001 | 0.21 | 0.0063 | 0.53 | 0.47 | | | | |
| | Gly | | 0.78 | 0.39 | 0.36 | 0.0001 | 0.0001 | 0.2 | 0.88 | 0.0007 | 0.57 | 0.016 | | | | |
| | Lys | **4** | 0.75 | 0.52 | 0.51 | 0.0001 | 0.0001 | 0.95 | 0.71 | 0.023 | 0.0087 | 0.0074 | 0.0001 | 0.041 | 0.0003 | 0.092 |
| | Ser | | 0.70 | 0.27 | 0.41 | 0.049 | 0.0001 | 0.0001 | 0.94 | 0.14 | 0.44 | 0.096 | | | | |
| | alpha-AAA | | 0.47 | 0.28 | 0.74 | 0.0001 | 0.017 | 0.39 | 0.92 | 0.67 | 0.14 | 0.011 | | | | |
| | Val | | 0.91 | 0.39 | 0.15 | 0.35 | 0.0001 | 0.0001 | 0.55 | 0.0009 | 0.2 | 0.027 | | | | |
| | Ile | **5** | 0.83 | 0.28 | 0.23 | 0.11 | 0.0001 | 0.011 | 0.63 | 0.0001 | 0.54 | 0.41 | 0.78 | 0.0001 | 0.0001 | 0.0002 |
| | Arg | | 0.67 | 0.46 | 0.62 | 0.074 | 0.0001 | 0.0012 | 0.75 | 0.0006 | 0.16 | 0.016 | | | | |
| Biogenic Amines | Spermidine | | 0.93 | 0.07 | 0.07 | 0.0011 | 0.012 | 0.88 | 0.69 | 0.092 | 0.12 | 0.97 | | | | |
| | Spermine | **1** | 0.88 | 0.01 | 0.12 | 0.25 | 0.064 | 0.77 | 0.27 | 0.2 | 0.12 | 0.83 | 0.0012 | 0.95 | 0.001 | 0.082 |
| | Putrescine | | 0.68 | 0.10 | 0.36 | 0.0001 | 0.0001 | 0.7 | 0.024 | 0.077 | 0.0007 | 0.81 | | | | |
| | Taurine | | 0.77 | 0.01 | 0.23 | 0.0002 | 0.13 | 0.0001 | 0.42 | 0.91 | 0.078 | 0.32 | | | | |
| | Creatinine | **2** | 0.72 | 0.01 | 0.29 | 0.029 | 0.067 | 0.012 | 0.62 | 0.032 | 0.15 | 0.75 | 0.0001 | 0.0001 | 0.049 | 0.23 |
| | Serotonin | | 0.49 | 0.00 | 0.51 | 0.0001 | 0.069 | 0.61 | 0.12 | 0.22 | 0.26 | 0.32 | | | | |
| | Carnosine | **3** | 0.79 | 0.04 | 0.22 | 0.0001 | 0.35 | 0.0005 | 0.072 | 0.69 | 0.12 | 0.38 | 0.0001 | 0.0001 | 0.55 | 0.21 |
| | SDMA | | 0.79 | 0.05 | 0.22 | 0.0001 | 0.94 | 0.0061 | 0.97 | 0.057 | 0.0018 | 0.39 | | | | |
| | C10:2 | | 0.85 | 0.47 | 0.28 | 0.0046 | 0.36 | 0.2 | 0.76 | 0.49 | 0.33 | 0.52 | | | | |
| | C14:2 | | 0.85 | 0.70 | 0.51 | 0.45 | 0.22 | 0.25 | 0.36 | 0.39 | 0.069 | 0.71 | | | | |
| | C14:2-OH | | 0.89 | 0.69 | 0.35 | 0.34 | 0.14 | 0.2 | 0.55 | 0.57 | 0.11 | 0.7 | | | | |
| | C16:2 | | 0.83 | 0.79 | 0.80 | 0.72 | 0.52 | 0.14 | 0.34 | 0.54 | 0.36 | 0.66 | | | | |
| | C3:1 | | 0.92 | 0.42 | 0.14 | 0.0005 | 0.13 | 0.6 | 1 | 0.6 | 0.075 | 0.54 | | | | |
| | C3-OH | | 0.92 | 0.44 | 0.15 | 0.0086 | 0.068 | 0.4 | 0.99 | 0.57 | 0.1 | 0.51 | | | | |
| | C4:1 | | 0.90 | 0.44 | 0.19 | 0.016 | 0.15 | 0.83 | 0.97 | 0.45 | 0.13 | 0.46 | | | | |
| | C5:1 | | 0.94 | 0.46 | 0.12 | 0.0013 | 0.096 | 0.47 | 0.99 | 0.55 | 0.24 | 0.52 | | | | |
| | C5:1-DC | **1** | 0.94 | 0.45 | 0.11 | 0.0003 | 0.092 | 0.53 | 0.96 | 0.7 | 0.11 | 0.41 | 0.016 | 0.32 | 0.2 | 0.54 |
| | C5-DC (C6-OH) | | 0.80 | 0.55 | 0.45 | 0.071 | 0.2 | 0.61 | 0.69 | 0.18 | 0.3 | 0.46 | | | | |
| | C5-M-DC | | 0.90 | 0.44 | 0.18 | 0.0027 | 0.039 | 0.61 | 0.98 | 0.37 | 0.1 | 0.42 | | | | |
| | C6 (C4:1-DC) | | 0.84 | 0.56 | 0.36 | 0.19 | 0.51 | 0.1 | 0.84 | 0.78 | 0.19 | 0.32 | | | | |
| | C6:1 | | 0.94 | 0.46 | 0.10 | 0.0027 | 0.34 | 0.35 | 0.96 | 0.32 | 0.27 | 0.53 | | | | |
| | C7-DC | | 0.89 | 0.57 | 0.26 | 0.072 | 0.22 | 0.07 | 0.57 | 0.13 | 0.25 | 0.63 | | | | |
| | C9 | | 0.91 | 0.53 | 0.20 | 0.0034 | 0.28 | 0.13 | 0.33 | 0.54 | 0.13 | 0.53 | | | | |
| | Sum of Medium-Chain ACs | | 0.88 | 0.54 | 0.26 | 0.011 | 0.31 | 0.15 | 0.74 | 0.82 | 0.15 | 0.33 | | | | |
| | C12 | | 0.85 | 0.53 | 0.32 | 0.0001 | 0.0004 | 0.0076 | 0.019 | 0.61 | 0.36 | 0.48 | | | | |
| | C14 | | 0.92 | 0.43 | 0.14 | 0.0001 | 0.0001 | 0.15 | 0.079 | 0.12 | 0.43 | 0.17 | | | | |
| | C16 | **2** | 0.94 | 0.51 | 0.12 | 0.0001 | 0.0001 | 0.24 | 0.16 | 0.062 | 0.65 | 0.38 | 0.0001 | 0.69 | 0.0001 | 0.48 |
| | C16:1 | | 0.93 | 0.55 | 0.16 | 0.0001 | 0.0001 | 0.054 | 0.0096 | 0.69 | 0.42 | 0.16 | | | | |
| | C18 | | 0.62 | 0.35 | 0.58 | 0.0001 | 0.077 | 0.01 | 0.9 | 0.0011 | 0.33 | 0.67 | | | | |
| | C18:1 | | 0.97 | 0.42 | 0.06 | 0.0001 | 0.0001 | 0.64 | 0.085 | 0.29 | 0.74 | 0.17 | | | | |

| | | | | | | | | | | | | | | | | |
|---|---|---|---|---|---|---|---|---|---|---|---|---|---|---|---|---|
| **Acyl carnitines** | C14:1 | **3** | 0.78 | 0.62 | 0.58 | 0.0001 | 0.024 | 0.043 | 0.021 | 0.75 | 0.21 | 0.47 | 0.022 | 0.12 | 0.039 | 0.54 |
| | C14:1-OH | | 0.92 | 0.68 | 0.26 | 0.11 | 0.015 | 0.074 | 0.29 | 0.33 | 0.29 | 0.34 | | | | |
| | C16:1-OH | | 0.91 | 0.56 | 0.21 | 0.0003 | 0.0027 | 0.28 | 0.31 | 0.47 | 0.31 | 0.15 | | | | |
| | C16:2-OH | | 0.87 | 0.70 | 0.42 | 0.1 | 0.023 | 0.18 | 0.29 | 0.41 | 0.18 | 0.55 | | | | |
| | C16-OH | | 0.83 | 0.53 | 0.36 | 0.074 | 0.086 | 0.16 | 0.048 | 0.046 | 0.11 | 0.85 | | | | |
| | C18:1-OH | | 0.83 | 0.58 | 0.41 | 0.57 | 0.4 | 0.15 | 0.018 | 0.73 | 0.14 | 0.57 | | | | |
| | C18:2 | | 0.63 | 0.56 | 0.85 | 0.0001 | 0.0049 | 0.29 | 0.22 | 0.23 | 0.74 | 0.36 | | | | |
| | Sum of ACs | | 0.91 | 0.65 | 0.25 | 0.41 | 0.012 | 0.11 | 0.16 | 0.85 | 0.35 | 0.14 | | | | |
| | Sum of Long-Chain ACs | | 0.91 | 0.63 | 0.25 | 0.0001 | 0.0081 | 0.16 | 0.15 | 0.32 | 0.6 | 0.26 | | | | |
| | Sum of MUFA-ACs | | 0.92 | 0.68 | 0.26 | 0.42 | 0.031 | 0.21 | 0.24 | 0.45 | 0.29 | 0.26 | | | | |
| | Sum of PUFA-ACs | | 0.86 | 0.57 | 0.33 | 0.37 | 0.095 | 0.12 | 0.41 | 0.54 | 0.43 | 0.35 | | | | |
| | 2MBG (NBS) | **4** | 0.75 | 0.21 | 0.32 | 0.0001 | 0.0001 | 0.26 | 0.82 | 0.0001 | 0.33 | 0.33 | 0.0001 | 0.17 | 0.0001 | 0.0001 |
| | C5 | | 0.67 | 0.35 | 0.51 | 0.0001 | 0.0001 | 0.86 | 0.27 | 0.0001 | 0.6 | 0.039 | | | | |
| | IVA (NBS) | | 0.81 | 0.21 | 0.24 | 0.0001 | 0.0001 | 0.17 | 0.35 | 0.0001 | 0.52 | 0.97 | | | | |
| | SBCAD Deficiency (NBS) | | 0.79 | 0.36 | 0.32 | 0.0001 | 0.0001 | 0.076 | 0.33 | 0.0001 | 0.86 | 0.32 | | | | |
| | SCAD Deficiency (NBS) | | 0.24 | 0.07 | 0.82 | 0.0001 | 0.63 | 0.85 | 0.3 | 0.16 | 0.075 | 0.28 | | | | |
| | 3MGA (NBS) | **5** | 0.65 | 0.09 | 0.38 | 0.0067 | 0.22 | 0.18 | 0.44 | 0.9 | 0.4 | 0.36 | 0.0001 | 0.022 | 0.032 | 0.41 |
| | BKT Deficiency (NBS) | | 0.57 | 0.11 | 0.48 | 0.0004 | 0.001 | 0.41 | 0.21 | 0.019 | 0.15 | 0.24 | | | | |
| | b-Oxidation | | 0.70 | 0.33 | 0.45 | 0.0004 | 0.21 | 0.12 | 0.44 | 0.062 | 0.84 | 0.14 | | | | |
| | CPT-1 Deficiency (NBS) | | 0.44 | 0.39 | 0.91 | 0.0001 | 0.54 | 0.0001 | 0.0011 | 0.0001 | 0.12 | 0.022 | | | | |
| | Ratio of Acetylcarnitine to Carnitine | | 0.65 | 0.06 | 0.38 | 0.0008 | 0.0099 | 0.082 | 0.69 | 0.0008 | 0.13 | 0.1 | | | | |
| | IBD Deficiency (NBS) | **6** | 0.78 | 0.19 | 0.27 | 0.0001 | 0.031 | 0.076 | 0.057 | 0.0001 | 0.14 | 0.95 | 0.0007 | 0.15 | 0.013 | 0.0014 |
| | MA (NBS) | | 0.89 | 0.17 | 0.13 | 0.085 | 0.0016 | 0.27 | 0.021 | 0.0051 | 0.0078 | 0.57 | | | | |
| | MMA (NBS) | | 0.73 | 0.35 | 0.42 | 0.0022 | 0.098 | 0.3 | 0.14 | 0.05 | 0.13 | 0.41 | | | | |
| | C0 | **7** | 0.74 | 0.38 | 0.43 | 0.0058 | 0.0009 | 0.013 | 0.054 | 0.0019 | 0.99 | 0.089 | 0.58 | 0.021 | 0.0008 | 0.003 |
| | C2 | | 0.67 | 0.23 | 0.43 | 0.68 | 0.0001 | 0.075 | 0.0027 | 0.0001 | 0.37 | 0.16 | | | | |
| | C3-DC (C4-OH) | | 0.57 | 0.31 | 0.62 | 0.047 | 0.03 | 0.0027 | 0.0001 | 0.0001 | 0.45 | 0.36 | | | | |
| | C5-OH (C3-DC-M) | | 0.79 | 0.49 | 0.41 | 0.086 | 0.007 | 0.14 | 0.085 | 0.044 | 0.41 | 0.042 | | | | |
| | Sum of Short-Chain Acs | | 0.79 | 0.66 | 0.63 | 0.14 | 0.0079 | 0.11 | 0.15 | 0.52 | 0.31 | 0.051 | | | | |
| | CACT Deficiency (NBS) | **8** | 0.59 | 0.40 | 0.69 | 0.0001 | 0.0049 | 0.0053 | 0.71 | 0.0001 | 0.98 | 0.67 | 0.0001 | 0.0029 | 0.36 | 0.024 |
| | LCHAD Deficiency (NBS) | | 0.74 | 0.32 | 0.39 | 0.0001 | 0.089 | 0.048 | 0.8 | 0.052 | 0.0019 | 0.097 | | | | |
| | Ratio of Medium-Chain to Long-Chain ACs | | 0.82 | 0.49 | 0.36 | 0.0001 | 0.022 | 0.087 | 0.074 | 0.17 | 0.62 | 0.97 | | | | |
| | VLCAD Deficiency (NBS) | | 0.65 | 0.36 | 0.54 | 0.0001 | 0.16 | 0.0006 | 0.34 | 0.014 | 0.063 | 0.25 | | | | |
| | w-Oxidation | | 0.53 | 0.30 | 0.67 | 0.0001 | 0.089 | 0.6 | 0.69 | 0.0096 | 0.31 | 0.76 | | | | |

| | | | | | | | | | | | | | | | | |
|---|---|---|---|---|---|---|---|---|---|---|---|---|---|---|---|---|
| | Ratio of Short-Chain to Medium-Chain ACs | 9 | 0.85 | 0.22 | 0.19 | 0.0029 | 0.14 | 0.5 | 0.92 | 0.093 | 0.64 | 0.63 | 0.0001 | 0.089 | 0.06 | 0.15 |
| | TFP Deficiency (NBS) | | 0.85 | 0.45 | 0.27 | 0.0001 | 0.051 | 0.0097 | 0.046 | 0.23 | 0.53 | 0.87 | | | | |
| | CPT-2 Deficiency (NBS) | 10 | 0.58 | 0.27 | 0.57 | 0.0001 | 0.12 | 0.001 | 0.097 | 0.0001 | 0.13 | 0.51 | 0.0001 | 0.0073 | 0.33 | 0.0001 |
| | MC Deficiency (NBS) | | 0.60 | 0.27 | 0.56 | 0.0001 | 0.03 | 0.089 | 0.41 | 0.0009 | 0.28 | 0.084 | | | | |
| | PA (NBS) | | 0.66 | 0.29 | 0.47 | 0.0001 | 0.063 | 0.029 | 0.054 | 0.0013 | 0.28 | 0.23 | | | | |
| | Ratio of Short-Chain to Long-Chain ACs | | 0.67 | 0.43 | 0.58 | 0.0001 | 0.62 | 0.81 | 0.88 | 0.0033 | 0.61 | 0.66 | | | | |
| | C3 | 11 | 0.91 | 0.23 | 0.11 | 0.21 | 0.61 | 0.2 | 0.84 | 0.012 | 0.58 | 0.048 | 0.0028 | 0.26 | 0.41 | 0.0039 |
| | C4 | | 0.91 | 0.30 | 0.12 | 0.0001 | 0.3 | 0.36 | 0.36 | 0.0024 | 0.4 | 0.05 | | | | |
| osphatidylcholines | PC aa C36:4 | 1 | 0.98 | 0.60 | 0.05 | 0.011 | 0.61 | 0.57 | 0.53 | 0.68 | 0.34 | 0.85 | 0.12 | 0.47 | 0.56 | 0.8 |
| | PC aa C32:0 | | 0.99 | 0.61 | 0.02 | 0.15 | 0.41 | 0.44 | 0.38 | 0.74 | 0.36 | 0.91 | | | | |
| | PC aa C38:4 | | 0.98 | 0.61 | 0.05 | 0.02 | 0.6 | 0.67 | 0.39 | 0.75 | 0.36 | 0.89 | | | | |
| | PC aa C32:1 | | 0.92 | 0.53 | 0.18 | 0.091 | 0.94 | 0.19 | 0.44 | 0.76 | 0.14 | 0.73 | | | | |
| | PC aa C36:1 | | 0.99 | 0.57 | 0.02 | 0.15 | 0.52 | 0.46 | 0.4 | 0.77 | 0.39 | 0.86 | | | | |
| | PC aa C38:6 | | 0.98 | 0.60 | 0.04 | 0.16 | 0.62 | 0.25 | 0.28 | 0.77 | 0.32 | 0.98 | | | | |
| | PC aa C40:6 | | 0.92 | 0.59 | 0.20 | 0.016 | 0.82 | 0.37 | 0.22 | 0.79 | 0.33 | 0.97 | | | | |
| | PC aa C34:1 | | 0.97 | 0.55 | 0.07 | 0.16 | 0.48 | 0.98 | 0.38 | 0.84 | 0.25 | 0.77 | | | | |
| | PC aa C36:2 | | 0.90 | 0.46 | 0.19 | 0.28 | 0.048 | 0.59 | 0.2 | 0.91 | 0.38 | 0.81 | | | | |
| | PC ae C42:2 | 2 | 0.84 | 0.60 | 0.39 | 0.0001 | 0.0008 | 0.0001 | 0.0041 | 0.036 | 0.047 | 0.14 | 0.0001 | 0.0001 | 0.0001 | 0.71 |
| | PC ae C36:3 | | 0.93 | 0.49 | 0.15 | 0.0001 | 0.2 | 0.0001 | 0.29 | 0.13 | 0.78 | 0.15 | | | | |
| | PC ae C38:3 | | 0.95 | 0.58 | 0.13 | 0.0001 | 0.011 | 0.0001 | 0.093 | 0.18 | 0.97 | 0.14 | | | | |
| | PC ae C38:1 | | 0.92 | 0.74 | 0.30 | 0.0001 | 0.0001 | 0.019 | 0.02 | 0.58 | 0.56 | 0.13 | | | | |
| | PC ae C38:2 | | 0.92 | 0.77 | 0.33 | 0.0001 | 0.0001 | 0.12 | 0.43 | 0.64 | 0.79 | 0.41 | | | | |
| | PC ae C40:2 | | 0.94 | 0.64 | 0.15 | 0.0001 | 0.0001 | 0.0001 | 0.0001 | 0.66 | 0.22 | 0.08 | | | | |
| | PC aa C42:6 | 3 | 0.82 | 0.41 | 0.31 | 0.0001 | 0.042 | 0.88 | 0.65 | 0.13 | 0.14 | 1 | 0.0001 | 0.19 | 0.76 | 0.84 |
| | PC aa C38:3 | | 0.82 | 0.58 | 0.43 | 0.0001 | 0.057 | 0.79 | 0.67 | 0.2 | 0.72 | 0.22 | | | | |
| | PC aa C40:4 | | 0.85 | 0.41 | 0.25 | 0.0001 | 0.34 | 0.0027 | 0.98 | 0.34 | 0.35 | 0.79 | | | | |
| | PC aa C42:4 | | 0.79 | 0.56 | 0.49 | 0.0001 | 0.81 | 0.79 | 0.23 | 0.49 | 0.098 | 0.89 | | | | |
| | PC aa C42:5 | | 0.81 | 0.54 | 0.40 | 0.0001 | 0.33 | 0.95 | 0.24 | 0.79 | 0.12 | 0.14 | | | | |
| | PC aa C38:5 | | 0.71 | 0.57 | 0.67 | 0.0001 | 0.84 | 0.97 | 0.78 | 0.79 | 0.28 | 0.3 | | | | |
| | PC aa C40:5 | | 0.54 | 0.33 | 0.70 | 0.0001 | 0.035 | 0.64 | 0.77 | 0.85 | 0.32 | 0.91 | | | | |
| | PC ae C36:5 | | 0.72 | 0.40 | 0.47 | 0.0001 | 0.71 | 0.0001 | 0.76 | 0.91 | 0.83 | 0.42 | | | | |
| | PC aa C26:0 | 4 | 0.88 | 0.44 | 0.22 | 0.077 | 0.43 | 0.93 | 0.11 | 0.067 | 0.94 | 0.97 | 0.96 | 0.55 | 0.38 | 0.27 |
| | PC aa C24:0 | | 0.85 | 0.59 | 0.36 | 0.092 | 0.61 | 0.62 | 0.22 | 0.088 | 0.92 | 0.94 | | | | |
| | PC aa C28:1 | | 0.78 | 0.57 | 0.51 | 0.0004 | 0.02 | 0.3 | 0.78 | 0.11 | 0.6 | 0.69 | | | | |
| | lysoPC a C24:0 | | 0.77 | 0.51 | 0.46 | 0.0088 | 0.45 | 0.27 | 0.81 | 0.54 | 0.83 | 0.81 | | | | |
| | lysoPC a C26:0 | | 0.89 | 0.38 | 0.18 | 0.99 | 0.83 | 0.88 | 0.085 | 0.55 | 0.97 | 0.74 | | | | |
| | lysoPC a C28:0 | | 0.88 | 0.40 | 0.19 | 0.83 | 0.39 | 0.11 | 0.12 | 0.67 | 0.98 | 0.92 | | | | |
| | lysoPC a C18:2 | 5 | 0.60 | 0.37 | 0.64 | 0.036 | 0.12 | 0.084 | 0.29 | 0.03 | 0.21 | 0.38 | 0.0003 | 0.074 | 0.0007 | 0.15 |
| | lysoPC a C18:0 | | 0.76 | 0.36 | 0.37 | 0.0001 | 0.0001 | 0.38 | 0.93 | 0.13 | 0.75 | 0.046 | | | | |
| | lysoPC a C16:1 | | 0.80 | 0.69 | 0.65 | 0.0007 | 0.02 | 0.0013 | 0.21 | 0.19 | 0.18 | 0.4 | | | | |
| | lysoPC a C17:0 | | 0.80 | 0.38 | 0.33 | 0.081 | 0.0003 | 0.054 | 0.56 | 0.33 | 0.36 | 0.12 | | | | |
| | PC aa C42:0 | 6 | 0.48 | 0.18 | 0.64 | 0.0001 | 0.11 | 0.34 | 0.46 | 0.18 | 0.065 | 0.61 | 0.0001 | 0.0001 | 0.0001 | 0.79 |
| | PC ae C42:1 | | 0.87 | 0.23 | 0.16 | 0.0001 | 0.0001 | 0.0001 | 0.027 | 0.29 | 0.0018 | 0.63 | | | | |
| | PC aa C42:2 | | 0.76 | 0.35 | 0.37 | 0.0001 | 0.0002 | 0.0001 | 0.19 | 0.38 | 0.11 | 0.33 | | | | |
| | PC aa C42:1 | | 0.80 | 0.25 | 0.26 | 0.0001 | 0.0087 | 0.0001 | 0.52 | 0.4 | 0.025 | 0.29 | | | | |
| | PC aa C40:1 | | 0.93 | 0.25 | 0.10 | 0.0001 | 0.0001 | 0.0001 | 0.05 | 0.67 | 0.046 | 0.34 | | | | |
| | PC aa C38:0 | | 0.71 | 0.38 | 0.47 | 0.0001 | 0.0001 | 0.031 | 0.028 | 0.67 | 0.46 | 0.64 | | | | |
| | PC aa C36:5 | 7 | 0.83 | 0.25 | 0.23 | 0.0001 | 0.0001 | 0.0001 | 0.2 | 0.0001 | 0.22 | 0.14 | 0.0001 | 0.0001 | 0.022 | 0.0033 |
| | PC aa C36:3 | | 0.77 | 0.47 | 0.44 | 0.0001 | 0.0072 | 0.0005 | 0.53 | 0.0008 | 0.67 | 0.37 | | | | |
| | PC aa C34:4 | | 0.78 | 0.38 | 0.36 | 0.14 | 0.16 | 0.0004 | 0.23 | 0.026 | 0.13 | 0.048 | | | | |
| | PC ae C42:3 | | 0.76 | 0.40 | 0.40 | 0.0001 | 0.72 | 0.0045 | 0.31 | 0.28 | 0.56 | 0.31 | | | | |
| | PC aa C40:3 | | 0.60 | 0.46 | 0.75 | 0.001 | 0.11 | 0.0001 | 0.0001 | 0.0002 | 0.66 | 0.062 | | | | |
| | PC ae C44:4 | | 0.50 | 0.27 | 0.68 | 0.0001 | 0.12 | 0.0001 | 0.019 | 0.069 | 0.1 | 0.57 | | | | |

| | | | | | | | | | | | | | | | | |
|---|---|---|---|---|---|---|---|---|---|---|---|---|---|---|---|---|
| Pl | PC ae C38:0 | 8 | 0.66 | 0.36 | 0.53 | 0.0001 | 0.25 | 0.0001 | 0.052 | 0.1 | 0.39 | 0.89 | 0.0001 | 0.0001 | 0.08 | 0.67 |
| | PC ae C40:1 | | 0.67 | 0.51 | 0.68 | 0.0001 | 0.0001 | 0.0001 | 0.1 | 0.52 | 0.52 | 0.79 | | | | |
| | PC ae C40:6 | | 0.52 | 0.33 | 0.72 | 0.1 | 0.037 | 0.0005 | 0.28 | 0.54 | 0.55 | 0.4 | | | | |
| | PC ae C44:6 | | 0.41 | 0.28 | 0.82 | 0.94 | 0.2 | 0.0001 | 0.011 | 0.56 | 0.0053 | 0.72 | | | | |
| | PC ae C44:5 | | 0.73 | 0.40 | 0.45 | 0.0001 | 0.85 | 0.0002 | 0.18 | 0.68 | 0.75 | 0.55 | | | | |
| | PC ae C42:0 | | 0.42 | 0.25 | 0.76 | 0.57 | 0.024 | 0.82 | 0.54 | 0.72 | 0.35 | 0.95 | | | | |
| | PC aa C36:6 | 9 | 0.43 | 0.27 | 0.77 | 0.0033 | 0.0089 | 0.0001 | 0.32 | 0.013 | 0.044 | 0.92 | 0.0001 | 0.12 | 0.042 | 0.76 |
| | PC ae C30:1 | | 0.81 | 0.39 | 0.31 | 0.0001 | 0.022 | 0.11 | 0.65 | 0.37 | 0.33 | 0.96 | | | | |
| | PC ae C30:2 | | 0.78 | 0.50 | 0.44 | 0.0003 | 0.13 | 0.12 | 0.54 | 0.65 | 0.12 | 0.76 | | | | |
| | PC ae C36:0 | | 0.79 | 0.47 | 0.39 | 0.0008 | 0.0036 | 0.34 | 0.14 | 0.66 | 0.29 | 0.96 | | | | |
| | PC aa C30:0 | | 0.84 | 0.61 | 0.42 | 0.044 | 0.066 | 0.048 | 0.14 | 0.74 | 0.023 | 0.66 | | | | |
| | PC ae C32:1 | | 0.73 | 0.47 | 0.51 | 0.0001 | 0.25 | 0.16 | 0.85 | 0.78 | 0.09 | 0.85 | | | | |
| | PC ae C30:0 | | 0.92 | 0.61 | 0.21 | 0.001 | 0.02 | 0.15 | 0.32 | 0.83 | 0.023 | 0.47 | | | | |
| | PC ae C34:0 | | 0.80 | 0.39 | 0.32 | 0.0001 | 0.037 | 0.24 | 0.24 | 0.88 | 0.2 | 0.83 | | | | |
| | PC ae C40:5 | 10 | 0.64 | 0.41 | 0.61 | 0.0001 | 0.21 | 0.033 | 0.25 | 0.0001 | 0.63 | 0.029 | 0.0001 | 0.67 | 0.0014 | 0.51 |
| | PC ae C40:4 | | 0.72 | 0.36 | 0.44 | 0.014 | 0.0008 | 0.47 | 0.63 | 0.084 | 0.63 | 0.053 | | | | |
| | PC ae C34:1 | | 0.78 | 0.29 | 0.30 | 0.0001 | 0.0001 | 0.64 | 0.66 | 0.099 | 0.61 | 0.0011 | | | | |
| | PC ae C38:4 | | 0.81 | 0.35 | 0.29 | 0.0001 | 0.047 | 0.61 | 0.51 | 0.27 | 0.42 | 0.004 | | | | |
| | PC ae C44:3 | | 0.50 | 0.42 | 0.86 | 0.0001 | 0.0088 | 0.28 | 0.13 | 0.44 | 0.15 | 0.23 | | | | |
| | PC ae C36:4 | | 0.74 | 0.50 | 0.53 | 0.0001 | 0.033 | 0.001 | 0.02 | 0.67 | 0.99 | 0.011 | | | | |
| | PC ae C38:5 | | 0.67 | 0.26 | 0.45 | 0.0001 | 0.011 | 0.034 | 0.25 | 0.88 | 0.92 | 0.024 | | | | |
| | PC ae C34:3 | 11 | 0.68 | 0.29 | 0.45 | 0.0018 | 0.94 | 0.12 | 0.95 | 0.0068 | 0.1 | 0.84 | 0.98 | 0.39 | 0.6 | 0.2 |
| | PC ae C32:2 | | 0.60 | 0.34 | 0.60 | 0.0013 | 0.012 | 0.14 | 0.58 | 0.072 | 0.65 | 0.16 | | | | |
| | lysoPC a C28:1 | | 0.72 | 0.42 | 0.49 | 0.0001 | 0.46 | 0.93 | 0.36 | 0.39 | 0.12 | 0.9 | | | | |
| | PC aa C32:3 | | 0.76 | 0.52 | 0.50 | 0.3 | 0.8 | 0.45 | 0.55 | 0.47 | 0.99 | 0.98 | | | | |
| | lysoPC a C26:1 | | 0.75 | 0.43 | 0.44 | 0.0021 | 0.69 | 0.81 | 0.64 | 0.49 | 0.13 | 0.8 | | | | |
| | PC aa C34:3 | 12 | 0.87 | 0.23 | 0.16 | 0.0048 | 0.041 | 0.0001 | 0.77 | 0.0002 | 0.53 | 0.58 | 0.13 | 0.0001 | 0.25 | 0.0003 |
| | PC aa C34:2 | | 0.87 | 0.32 | 0.18 | 0.41 | 0.83 | 0.0001 | 0.35 | 0.0009 | 0.11 | 0.53 | | | | |
| | lysoPC a C18:1 | 13 | 0.84 | 0.63 | 0.44 | 0.0039 | 0.0003 | 0.4 | 0.065 | 0.02 | 0.52 | 0.31 | 0.52 | 0.15 | 0.0021 | 0.075 |
| | lysoPC a C20:3 | | 0.78 | 0.36 | 0.34 | 0.0002 | 0.062 | 0.0004 | 0.46 | 0.026 | 0.14 | 0.42 | | | | |
| | lysoPC a C20:4 | | 0.86 | 0.29 | 0.20 | 0.0009 | 0.001 | 0.74 | 0.049 | 0.045 | 0.51 | 0.51 | | | | |
| | lysoPC a C16:0 | | 0.91 | 0.76 | 0.36 | 0.66 | 0.001 | 0.32 | 0.32 | 0.15 | 0.25 | 0.29 | | | | |
| | PC ae C40:3 | 14 | 0.73 | 0.55 | 0.59 | 0.0001 | 0.033 | 0.0001 | 0.0001 | 0.0046 | 0.62 | 0.12 | 0.0001 | 0.0085 | 0.0001 | 0.27 |
| | PC ae C36:1 | | 0.75 | 0.53 | 0.52 | 0.0001 | 0.0001 | 0.0029 | 0.27 | 0.08 | 0.79 | 0.037 | | | | |
| | PC ae C34:2 | | 0.69 | 0.34 | 0.47 | 0.0001 | 0.0001 | 0.57 | 0.27 | 0.082 | 0.84 | 0.049 | | | | |
| | PC aa C36:0 | | 0.64 | 0.38 | 0.58 | 0.0001 | 0.0001 | 0.68 | 0.0062 | 0.18 | 0.18 | 0.59 | | | | |
| | PC ae C36:2 | | 0.88 | 0.79 | 0.59 | 0.0001 | 0.0008 | 0.2 | 0.85 | 0.31 | 0.86 | 0.16 | | | | |
| | PC aa C40:2 | | 0.74 | 0.51 | 0.53 | 0.0001 | 0.0001 | 0.0001 | 0.0047 | 0.56 | 0.42 | 0.28 | | | | |
| Sphingomyelins | SM C18:0 | 1 | 0.90 | 0.29 | 0.14 | 0.038 | 0.29 | 0.43 | 0.32 | 0.89 | 0.16 | 0.72 | 0.48 | 0.045 | 0.079 | 0.8 |
| | SM (OH) C14:1 | | 0.81 | 0.44 | 0.34 | 0.65 | 0.11 | 0.074 | 0.2 | 0.41 | 0.065 | 0.29 | | | | |
| | SM C20:2 | | 0.81 | 0.38 | 0.32 | 0.0063 | 0.8 | 0.32 | 0.43 | 0.79 | 0.049 | 0.7 | | | | |
| | SM (OH) C24:1 | | 0.77 | 0.29 | 0.32 | 0.081 | 0.095 | 0.004 | 0.2 | 0.13 | 0.21 | 0.68 | | | | |
| | SM C24:0 | | 0.73 | 0.57 | 0.64 | 0.71 | 0.0002 | 0.047 | 0.022 | 0.64 | 0.28 | 0.3 | | | | |
| | SM (OH) C16:1 | | 0.69 | 0.33 | 0.46 | 0.44 | 0.38 | 0.12 | 0.14 | 0.67 | 0.07 | 0.15 | | | | |
| | SM C16:0 | | 0.65 | 0.57 | 0.81 | 0.0001 | 0.18 | 0.088 | 0.046 | 0.71 | 0.17 | 0.2 | | | | |
| | SM C26:0 | | 0.58 | 0.31 | 0.61 | 0.0005 | 0.0001 | 0.0018 | 0.1 | 0.82 | 0.59 | 0.21 | | | | |
| | SM C24:1 | 2 | 0.88 | 0.21 | 0.15 | 0.0001 | 0.0001 | 0.0014 | 0.0003 | 0.11 | 0.62 | 0.097 | 0.0001 | 0.0057 | 0.0001 | 0.81 |
| | SM C26:1 | | 0.86 | 0.67 | 0.43 | 0.0001 | 0.0006 | 0.0098 | 0.023 | 0.95 | 0.2 | 0.35 | | | | |
| | SM (OH) C22:2 | | 0.80 | 0.15 | 0.24 | 0.0001 | 0.0001 | 0.0008 | 0.0001 | 0.0033 | 0.2 | 0.01 | | | | |
| | SM (OH) C22:1 | | 0.75 | 0.60 | 0.62 | 0.0001 | 0.033 | 0.059 | 0.27 | 0.76 | 0.084 | 0.36 | | | | |
| | SM C16:1 | 3 | 0.85 | 0.39 | 0.25 | 0.12 | 0.68 | 0.014 | 0.25 | 0.12 | 0.39 | 0.34 | 0.62 | 0.19 | 0.49 | 0.23 |
| | SM C18:1 | | 0.85 | 0.14 | 0.18 | 0.0007 | 0.093 | 0.96 | 0.67 | 0.37 | 0.5 | 0.72 | | | | |
