## Supplementary Table S3 for "APOE Genotype Influences on The Brain Metabolome of Aging Mice – Role for Mitochondrial Energetics in Mechanisms of Resilience in APOE2 Genotype"

**Table S3. APOE genotypes influenced multiple metabolism pathways in the brains in the ROS-MAP cohort.** ANCOVA analysis was performed on medication-adjusted metabolomic data, controlling BMI, PMI, age at death, education, sex, cognitive diagnosis, and beta-hydroxyisovaleroylcarnitine, followed by Tukey pos-hoc test, where the levels were not decorated with the same letters were significantly different from each other (i.e., A and B).

| Metabolite | APOE genotype | | | p-value | SUPER_PATHWAY | SUB_PATHWAY |
|---|---|---|---|---|---|---|
| | e2e3 | e3e3 | e3e4 | | | |
| carnitine | B | A | A | 0.0042 | Lipid | Carnitine Metabolism |
| (S)-3-hydroxybutyrylcarnitine | B | A | A | 0.0075 | Lipid | Fatty Acid Metabolism (Acyl Carnitine, Hydroxy) |
| eicosenoylcarnitine (C20:1)* | B | AB | A | 0.0412 | Lipid | Fatty Acid Metabolism (Acyl Carnitine, Monounsaturated) |
| arachidonoylcarnitine (C20:4) | B | AB | A | 0.0254 | Lipid | Fatty Acid Metabolism (Acyl Carnitine, Polyunsaturated) |
| acetylcarnitine (C2) | B | A | A | 0.0033 | Lipid | Fatty Acid Metabolism (Acyl Carnitine, Short Chain) |
| methylmalonate (MMA) | AB | A | B | 0.0049 | Lipid | Fatty Acid Metabolism (also BCAA Metabolism) |
| alpha-hydroxyisovalerate | B | A | AB | 0.0215 | Amino Acid | Leucine, Isoleucine and Valine Metabolism |
| tiglyl carnitine (C5) | B | A | AB | 0.0265 | Amino Acid | Leucine, Isoleucine and Valine Metabolism |
| 3-methylglutaconate | B | A | AB | 0.0471 | Lipid | Leucine, Isoleucine and Valine Metabolism |
| N-behenoyl-sphingadienine (d18:2/22:0)* | A | A | A | 0.0330 | Lipid | Ceramides |
| stearoyl-docosahexaenoyl-glycerol (18:0/22:6) [1]* | A | B | AB | 0.0348 | Lipid | Diacylglycerol |
| 1-stearoyl-GPC (18:0) | A | B | AB | 0.0038 | Lipid | Lysophospholipid |
| 1-palmitoyl-GPC (16:0) | A | A | A | 0.0490 | Lipid | Lysophospholipid |
| 1-oleoyl-2-docosahexaenoyl-GPC (18:1/22:6)* | A | A | A | 0.0290 | Amino Acid | Phosphatidylcholine (PC) |
| 1-palmitoyl-2-docosahexaenoyl-GPC (16:0/22:6) | AB | B | A | 0.0230 | Lipid | Phosphatidylcholine (PC) |
| N-acetylglucosamine/N-acetylgalactosamine | A | AB | B | 0.0480 | Carbohydrate | Aminosugar Metabolism |
| benzoate | A | AB | B | 0.0485 | Xenobiotics | Benzoate Metabolism |
| arachidonoyl ethanolamide | A | B | B | 0.0142 | Lipid | Endocannabinoid |
| methyl glucopyranoside (alpha + beta) | A | A | B | 0.0022 | Xenobiotics | Food Component/Plant |
| 3-hydroxystachydrine* | A | AB | B | 0.0071 | Xenobiotics | Food Component/Plant |
| stachydrine | A | B | B | 0.0087 | Xenobiotics | Food Component/Plant |
| ophthalmate | A | A | A | 0.0498 | Amino Acid | Glutathione Metabolism |
| imidazole propionate | AB | B | A | 0.0231 | Amino Acid | Histidine Metabolism |
| chiro-inositol | A | AB | B | 0.0368 | Lipid | Inositol Metabolism |
| S-adenosylhomocysteine (SAH) | B | A | AB | 0.0350 | Amino Acid | Methionine, Cysteine, SAM and Taurine Metabolism |
| 2,3-dihydroxy-5-methylthio-4-pentenoate (DMTPA)* | B | A | A | 0.0095 | Lipid | Methionine, Cysteine, SAM and Taurine Metabolism |
| 2'-deoxycytidine | A | AB | B | 0.0355 | Amino Acid | Pyrimidine Metabolism, Cytidine containing |
| cytidine | A | AB | B | 0.0440 | Nucleotide | Pyrimidine Metabolism, Cytidine containing |
| kynurenate | B | A | A | 0.0228 | Amino Acid | Tryptophan Metabolism |
| kynurenine | B | A | A | 0.0232 | Amino Acid | Tryptophan Metabolism |
| tyrosine | B | A | AB | 0.0408 | Amino Acid | Tyrosine Metabolism |
| pyridoxal | A | B | AB | 0.0431 | Cofactors and Vitamins | Vitamin B6 Metabolism |
| X - 24431 | A | B | AB | 0.0078 | 0 | 0 |
| X - 23171 | AB | B | A | 0.0312 | 0 | 0 |
| X - 24807 | A | B | AB | 0.0427 | 0 | 0 |
| X - 12104 | A | A | A | 0.0472 | 0 | 0 |
