## Supplementary Table S3 for "APOE Genotype Influences on The Brain Metabolome of Aging Mice – Role for Mitochondrial Energetics in Mechanisms of Resilience in APOE2 Genotype"

### Table S4. Cluster components of selected ROS-MAP metabolites.

| Compound | Cluster information | | | | P values - Full factorial model on cluster components | | | | | | | |
|---|---|---|---|---|---|---|---|---|---|---|---|---|
| | **Cluster** | **RSquare with Own Cluster** | **RSquare with Next Closest** | **1-RSquare Ratio** | **Age at death** | **BMI** | **APOE Genotypes** | **Beta-hydroxyisovaleroylcarnitine** | **Cognition diagnosis** | **Education** | **Sex** | **PMI** |
| (S)-3-hydroxybutyrylcarnitine | 1 | 0.626 | 0.072 | 0.402 | 0.0934 | 0.9953 | **0.0002** | **<.0001** | 0.9502 | 0.4869 | 0.2048 | **<.0001** |
| acetylcarnitine (C2) | | 0.635 | 0.106 | 0.409 | | | | | | | | |
| arachidonoylcarnitine (C20:4) | | 0.389 | 0.001 | 0.611 | | | | | | | | |
| carnitine | | 0.487 | 0.113 | 0.578 | | | | | | | | |
| eicosenoylcarnitine (C20:1)* | | 0.616 | 0.142 | 0.448 | | | | | | | | |
| tiglyl carnitine (C5:1) | | 0.552 | 0.266 | 0.61 | | | | | | | | |
| 3-methylglutaconate | 2 | 0.668 | 0.116 | 0.375 | 0.5889 | 0.2374 | **0.0023** | **<.0001** | 0.5082 | 0.1103 | 0.0573 | **0.0100** |
| alpha-hydroxyisovalerate | | 0.612 | 0.198 | 0.484 | | | | | | | | |
| methylmalonate (MMA) | | 0.396 | 0.019 | 0.615 | | | | | | | | |
